# Cryo-EM structure of DDM1-HELLS chimera bound to nucleosome reveals a mechanism of chromatin remodeling and disease regulation

**DOI:** 10.1101/2023.08.09.551721

**Authors:** Wilson Nartey, Aaron A. Goodarzi, Gareth J. Williams

## Abstract

Human HELicase, Lymphoid Specific (HELLS), and plant homolog Deficient in DNA Methylation 1 (DDM1), belong to a distinct class of chromatin remodelers that play important roles in DNA repair, transcription, and maintenance of DNA methylation in heterochromatin. HELLS also promotes the growth of hard-to-treat cancers including glioblastoma and hepatocellular carcinoma. Here, we identify an auto-inhibitory HELLS N-terminal coiled-coil, unravelling a long-standing question of HELLS inactivity *in vitro*. Using cryo-EM, we determine the 3.5 Å structure of an active DDM1-HELLS chimera in complex with a nucleosome. The structure reveals that a HELLS-specific insertion in the ATPase lobe 2 interacts with the nucleosome acidic patch to enhance chromatin remodeling. At the C-terminus, we resolve a unique motif, and disease hot spot, that binds and stabilizes the ATPase motor of the HELLS family of remodelers. Finally, we provide mechanistic insights for how post-translational modifications in the motor domain and midloop could modulate HELLS activity to regulate cancer stem cell state.

## Main

Chromatin is a barrier to DNA replication, transcription, and repair. To overcome this obstacle, ATP-dependent chromatin remodelers are recruited to regulate access to DNA by sliding or evicting nucleosomes^1,2^. The SNF2 family of chromatin remodelers, which includes SWI/SNF, ISWI, CHD, and INO80, contain RecA-like domains that utilize energy from ATP hydrolysis to move DNA over histone octamers^3,4^.

Human HELicase, Lymphoid Specific (HELLS), and plant Deficient in DNA Methylation 1 (DDM1) belong to a distinct class of SNF2 chromatin remodelers that are essential for maintaining genome-wide DNA methylation by facilitating access to DNA in heterochromatin^5–8^. HELLS mutations result in the DNA methylation associated disease Immunodeficiency-Centromeric instability-Facial abnormalities syndrome 4 (ICF4)^9–11^. Deletion, knockdown, or mutation of either HELLS or DDM1-encoding genes leads to genomic instability because of their roles in DNA damage repair in heterochromatin^9,12–17^. HELLS and DDM1 have also been shown to deposit histone variants, macroH2A and H2A.W respectively, into chromatin to repress expression of target genes and transposable elements^18–21^ and promote homologous recombination-mediated DNA double strand break repair at stalled replication forks^19^

Multifunctional roles of HELLS in cells are mediated by protein-protein and protein-RNA interactions^5,14,22–31^. The overexpression of HELLS promotes tumor proliferation, metastasis, and maintains cancer stem cell state^31–40^. In glioblastoma brain cancer cells, HELLS binds E2F3 and c-Myc to regulate the expression of target genes to maintain stemness^23^. The phosphorylation and methylation of HELLS also modulates the cancer cell stemness^41^, and overexpression also is linked with epigenetic silencing of tumor suppressors by increasing nucleosome occupancy at target genes in hepatocellular carcinoma^42^. Targeted inhibition of HELLS has, therefore, emerged as a viable therapeutic approach for high lethality cancers^23^.

Unlike DDM1, which has been shown to actively slide nucleosomes *in vitro*^43^, purified HELLS fails to remodel reconstituted nucleosomes, and has minimal ATPase activity^9,16,17^, although activities are enhanced when complexed with Cell Division Cycle Associated 7 (CDCA7)^9,10^. The lack of intrinsic nucleosome remodeling activity raises the possibility that HELLS is auto-inhibited, and this auto-inhibition is not overcome by the addition of substrate unlike other remodelers including CHD1 and ISWI^44–46^. However, the absence of HELLS and DDM1 structures, coupled with limited biochemical and mechanistic studies on this distinct class of SNF2 chromatin remodelers, has impeded our understanding of their biological functions and the design of effective cancer therapeutics targeting HELLS.

In this study, we compared the enzymatic activities of HELLS and DDM1 and discovered an autoinhibitory helical element located at the N-terminus of HELLS. Additionally, we identified a unique insertion in the HELLS motor domain that stimulates chromatin remodeling activity. Using cryo-Electron Microscopy (cryo-EM), we determined the 3.5 Å structure of a DDM1-HELLS chimeric protein bound to a nucleosome. This structure reveals interactions of the DDM1 motor with nucleosomal DNA and the histone H4 N-terminal tail, with mutation of key interacting residues reducing nucleosome remodeling *in vitro*. Furthermore, our structure indicates that the unique insertion in the HELLS motor domain interacts extensively with the nucleosome acidic patch. Mutational studies shed light on the role of conserved arginine residues and a ’tyrosine anchor’, as well as a proline-induced conformational constraint on acidic patch recognition. Our structure, along with molecular dynamic simulations, reveal mechanistic insights into the role of post-translational modifications in regulating chromatin remodeling and cancer cell stemness. Finally, we provide results and insights into how HELLS mutations found in cancer genomic databases or causing ICF4 syndrome impact HELLS activities and stability.

## Results

### An N-terminal coiled-coil domain autoinhibits HELLS

To understand the underlying mechanism of HELLS regulation, we first compared the biochemical activities of HELLS to its plant homolog, DDM1, and the human ISWI enzyme called SNF2H/SMARCA5 (Fig. 1). We cloned and expressed all proteins in *Escherichia coli* and purified them to homogeneity using a final gel filtration step (Extended Fig. 1a, 1b). Comparing the DNA and nucleosome-stimulated ATPase activity of HELLS and DDM1 under the same experimental conditions, we found that, despite good homology (39% identity, 54% similarity, Extended Data Fig. 1), HELLS exhibits minimal ATPase activity, whereas DDM1 hydrolyzes ATP robustly in the presence of substrates (Fig. 1b). We then measured the ATPase-dependent nucleosome remodeling activity using end-positioned nucleosomes in a bulk FRET assay^47^. Consistent with the ATPase measurements, HELLS was unable to remodel the nucleosome, whereas DDM1 and SNF2H had robust remodeling activity, with SNF2H having approximately 6-fold higher activity (Fig. 1c). These results are consistent with previous studies that compared DDM1 to SNF2H^43^ and confirm the possibility that HELLS is autoinhibited.

**Fig. 1.**
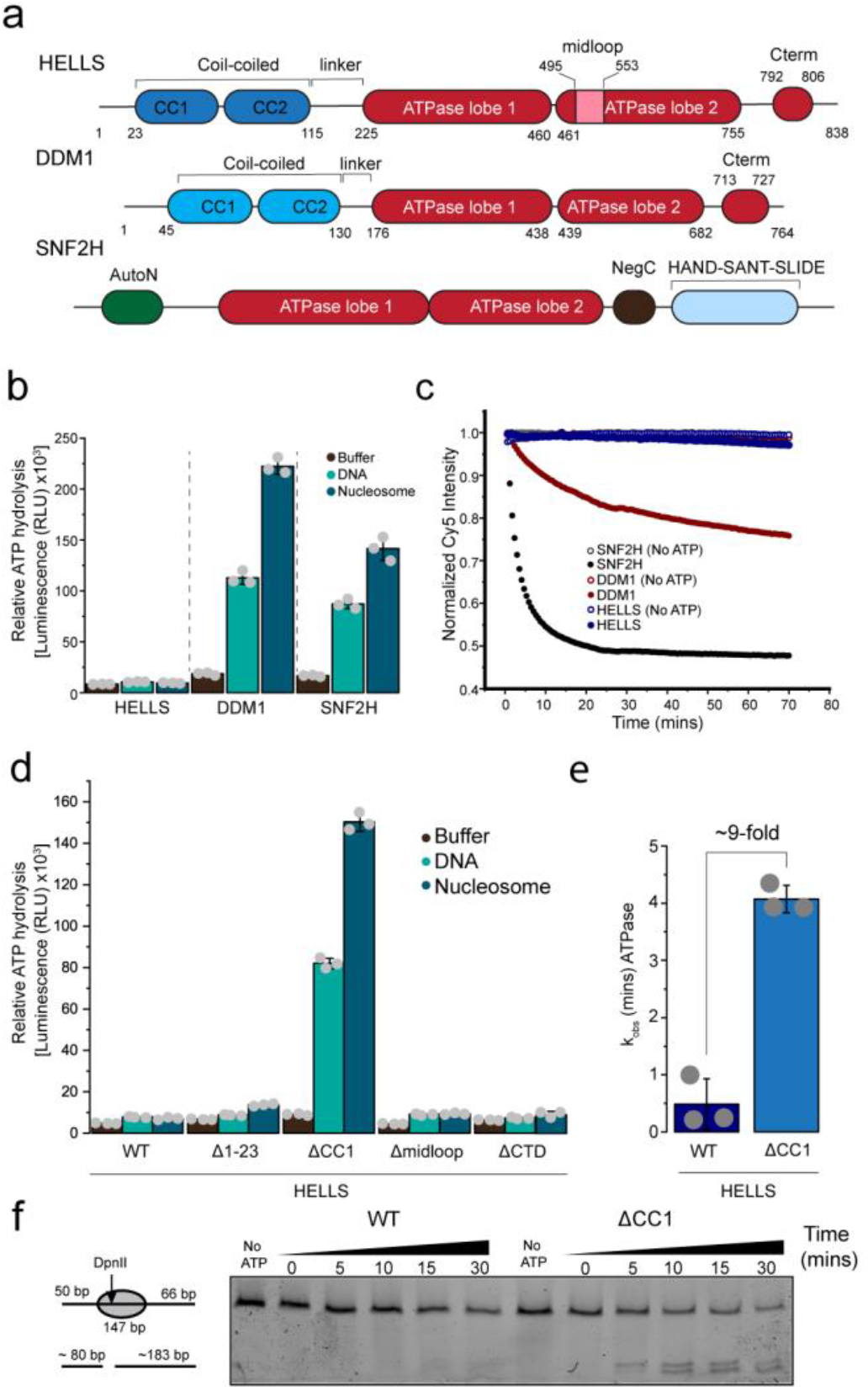
| HELLS is autoinhibited by the N-terminal coiled-coil domain. **a,** Domain achitecture of hsHELLS, atDDM1 and hsSNF2H. **b,** Endpoint ATP hydrolysis measurements of chromatin remodellors (0.2 µM) in the absence or precence of DNA (0.2 µM) and 30N30 nucleosome (0.2 µM). The error bars represent s.d. of three technical replicates. **c,** Average FRET traces (3 technical replicates) of chromatin remodeling activity of indicated proteins (1 μM) with 0.03 μM Cy3 and Cy5 labelled 0N50 nucleosome. hsSNF2H and atDDM1 traces were fiited with a double exponential and single exponetial equations, respectively. **d,** Endpoint ATP hydrolysis measurements for indicated HELLS variants containing deletion of internal motifs, measured as in **b. e,** Comparism of ATP hydrolysis rates between HELLS and ΔCC1-HELLS in the presence of 0.05 µM 30N30 nuclesosme. Data is mean ± s.d from three independent measurements. **f,** Representative gel of a restriction enzyme accessibility assay (5% TBE), demonstrating the activation of HELLS by CC1 deletion. HELLS proteins and 50N66 nucleosome (Epicypher) were used at 0.4 µM and 0.03 µM, respectively. Source data for **b**, **c**, **d** and **e** are provided.

Sequence analysis reveals unique features of HELLS and DDM1 compared to other SNF2 family members. At the N-terminus, HELLS and DDM1 contain predicted coiled-coil domains. Within the ATPase motor HELLS contains a novel 58 amino acid insertion that we call the ‘midloop’, which is absent in other chromatin remodelers, including DDM1. At the C-terminus, there is a conserved motif only present in the HELLS/DDM1 family of remodelers that we call the ‘Cterm’ motif (Extended Data Fig. 2).

**Fig. 2.**
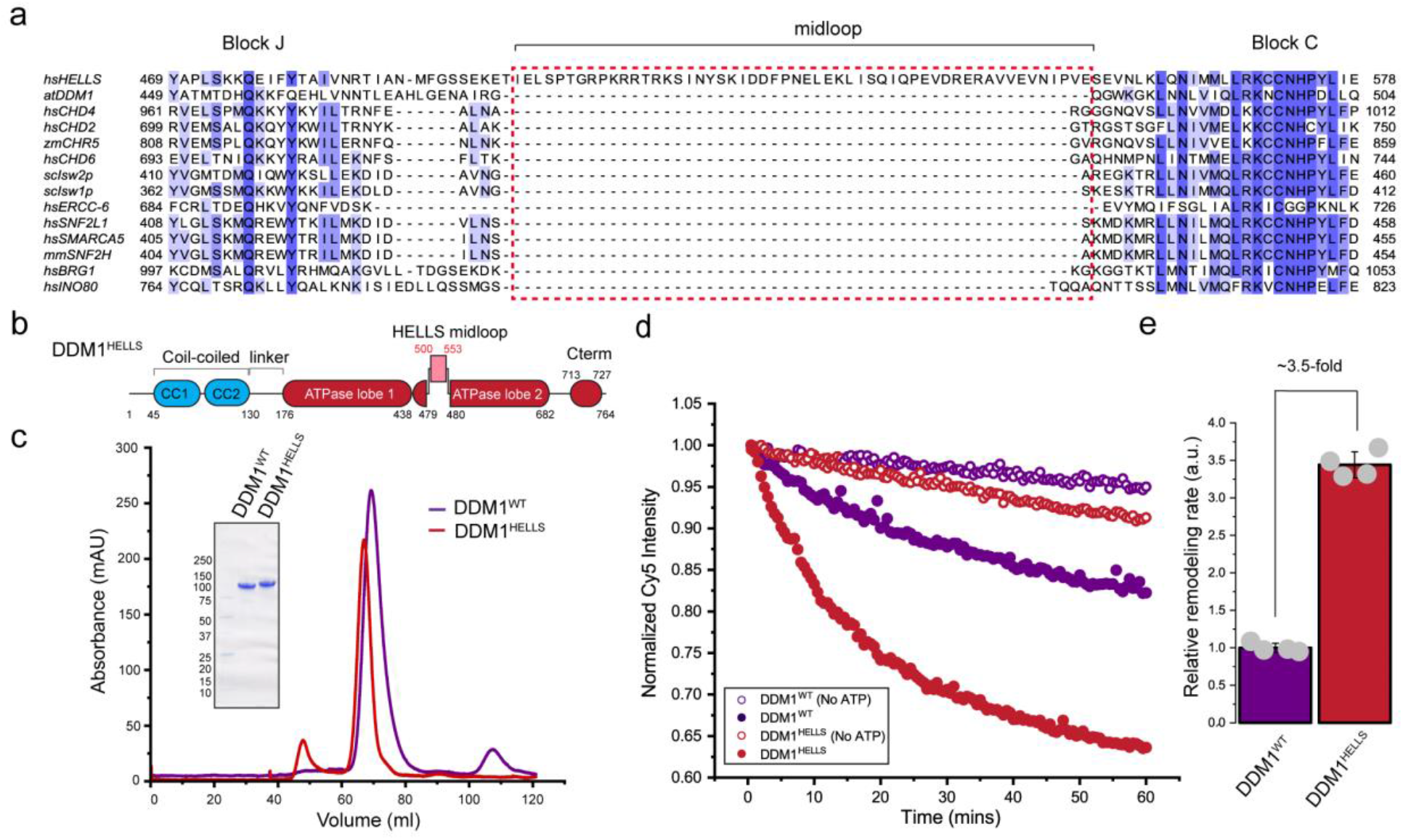
| The midloop of HELLS stimulates chromatin remodeling. **a,** Sequence alignment of the variable loop regions of Snf2 chromatin remodelers. The unique insertion of a midloop polypeptide in the HELLS sequence is highlighted (red box). **b,** Schematic representation of the generation of DDM1^HELLS^ chimera. **c,** Size-exclusion chromatography profile of DDM1 and DDM1^HELLS^ proteins on a Hiload 16/600 Superdex^TM^ 200 pg column. Coomassie-stained SDS-PAGE of purified proteins (inset). **d,** Representative FRET traces comparing DDM1 (1.0 µM) and DDM1^HELLS^ (1.0 µM) using 0.03 µM nucleosome substrate in the presence or absence of 2 mM ATP. **e,** Quantification of FRET traces after double (DDM1^HELLS^) or single (DDM1) exponential fit. Data is mean ± s.d from four independent experiments. Source data for **d** and **e** are provided.

To identify which region might be responsible for HELLS autoinhibition, we sequentially deleted the unique regions across HELLS and tested for ATPase activity (Fig. 1d, Extended Data Fig. 1d and e). An autoinhibitory domain was mapped to the N-terminal region of HELLS, as deletion of the first of two predicted coiled-coil helices (ΔCC1, amino acids A24-S55) robustly stimulated the ATPase activity of HELLS in the presence of DNA and nucleosome. To validate this observation, we produced an ATPase-dead HELLS mutant (ΔCC1-HELLS K254R) that abolishes ATPase activity (Extended Data Fig. 1f). We then performed an NADH-coupled ATPase assay revealing a 9-fold stimulation of the ΔCC1-HELLS mutant ATPase activity compared to wild-type HELLS, under single-turnover conditions (excess enzymes) in the presence of nucleosomes (Fig. 1e). Finally, we confirmed that the activation of ΔCC1-HELLS ATPase activity promotes chromatin remodeling using centrally positioned nucleosomes (Fig. 1f). Taken together, these results reveal a previously unknown autoinhibitory mechanism involving the N-terminal coiled-coil domain of HELLS.

### The midloop of HELLS stimulates chromatin remodeling

The existence of the midloop, a unique 58 amino acid insertion in lobe 2 of the ATPase motor of HELLS, prompted us to investigate its role in regulating HELLS activity (Fig. 2a, Extended Data Fig. 2). The region of the midloop insertion, known as HD2 (α-Helical Domain 2) or protrusion 2, is specific to the SNF2 family of remodelers and plays important roles^48–50^. It contains a ‘gating helix’ that contacts the tracking strand to orchestrate the forward movement of the translocating DNA^3,4^. A short variable loop, which in many remodelers is adjacent to the L1/L2 loops of histone H3, connects the gating helix and another helical component of HD2^4^ (Extended Data Fig. 3a). In HELLS, this short loop is replaced with the unusually large midloop (Fig. 2a, Extended Data Fig. 2). Using constitutively active DDM1 as a surrogate, we engineered the human HELLS midloop into the equivalent region of DDM1 (between residues G479 and Q480), to form a DDM1-HELLS chimeric protein (DDM1^HELLS^, Fig 2b). We subjected both DDM1 and DDM1^HELLS^ to size exclusion chromatography and mass photometry, confirming both proteins to be homogeneous and monomeric (Fig. 2c, Extended Data Fig. 3b). Biochemical comparisons show that while DDM1 and DDM1^HELLS^ bind DNA and nucleosomes with similar affinity (Extended Data Fig. 3c, d), the rate of ATP hydrolysis differs depending on the substrate. DNA stimulates both proteins to a similar extent, whereas nucleosomes stimulate DDM1^HELLS^ ATPase activity over 2-fold more than it did for DDM1 (Extended Data Fig. 3e). Furthermore, measuring nucleosome remodeling activity using our FRET assay with an end-positioned nucleosome reveals a 3.5-fold increase in DDM1^HELLS^ activity compared to DDM1 (Fig. 2d and e). Taken together, our results suggest that the HELLS midloop stimulates ATPase activity and nucleosome remodeling in a nucleosome-dependent manner, leading us to conduct structural studies of DDM1^HELLS^ bound to a nucleosome.

**Fig. 3.**
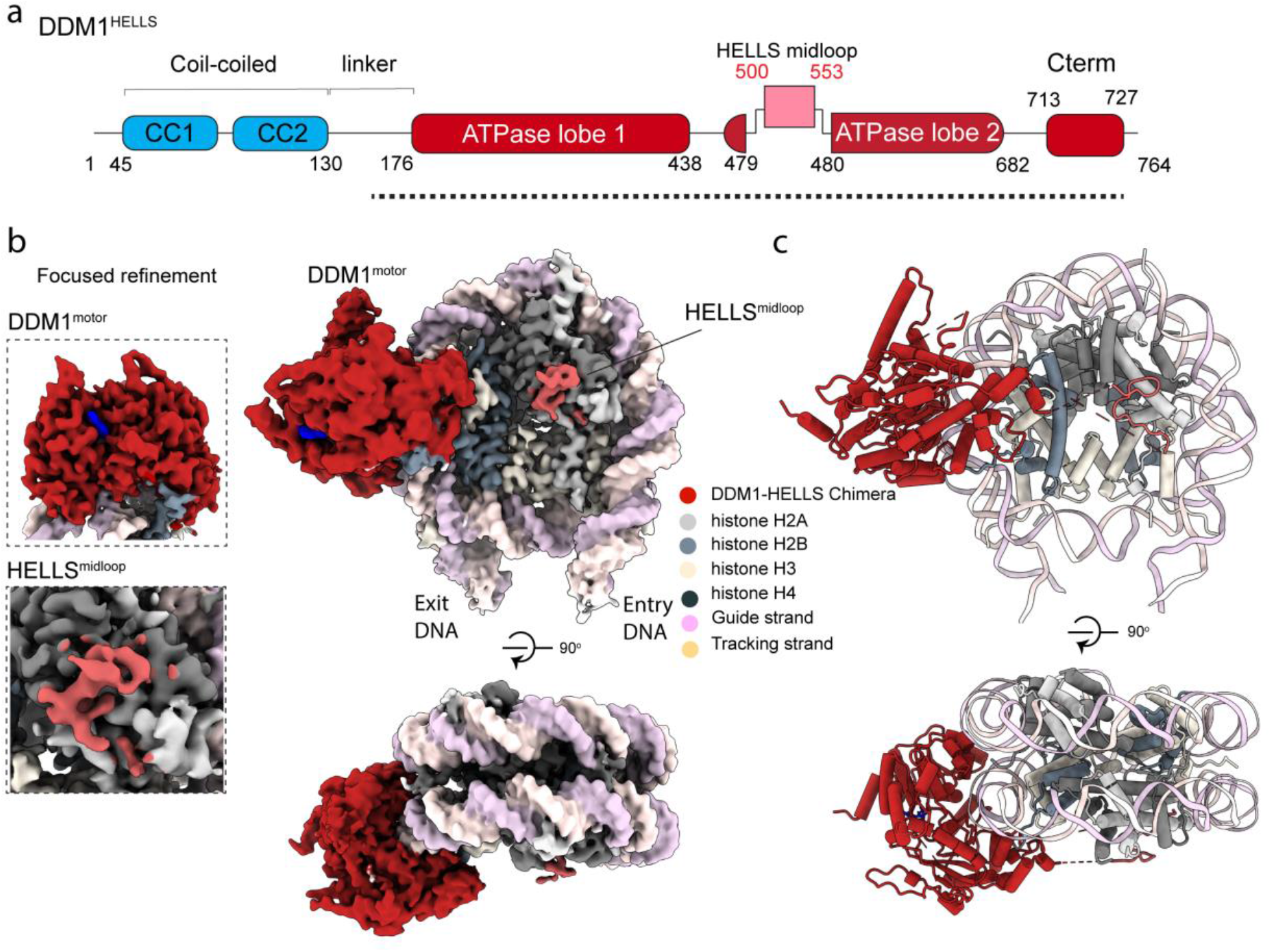
| Cryo-EM structure of DDM1^HELLS^ bound to the nucleosome. **a,** Structural features resolved in the cryo-EM reconstruction are marked by dotted line on the domain architecture of DDM1^HELLS^**. b,** Two views of reconstructed structures of DDM1^HELLS^ are shown (*right*), revealing interaction of DDM1 motor protein at SHL2 and the midloop of HELLS interaction with the H2A/H2B dimer of the nucleosome. Focused refinement of the DDM1 motor and HELLS midloop are shown as inset (*left*). **c,** Equivalent views as in **b,** of a cartoon representation of the DDM1^HELLS^- nucleosome complex model.

### Cryo-EM structure of DDM1^HELLS^ chimera bound to the nucleosome

To understand the structural basis of HELLS and DDM1 chromatin remodeling and evaluate the stimulatory contributions of the HELLS midloop, we determined the cryo-EM structure of DDM1^HELLS^ in complex with a nucleosome (Fig. 3a, Extended Data Fig. 4a-e, Supplementary Video 1, Table 1). Reconstituted nucleosomes, using Widom 601 DNA with 30 and 15 bases at the respective entry and exit sites (30N15), were mixed with DDM1^HELLS^ and ADP.BeF_x_, and the complex further stabilized by glutaraldehyde cross-linking. Following cryo-EM data collection and processing we resolved the 1:1 and 2:1 DDM1^HELLS^: nucleosome complexes at nominal resolution of 3.5 Å. We used focused refinement on the 1:1 DDM1^HELLS^-nucleosome complex to further improve the resolution of the DDM1 motor domain and HELLS midloop (Extended Data 4a-e). The level of detail observed in our reconstruction allowed us to build an atomic model of DDM1^HELLS^ bound to a mononucleosome (Fig. 3b, Extended Data Fig. 5). Although we resolved the 2:1 cryo-EM density of the complex, we did not proceed to build atomic models because analysis of the maps did not reveal any additional information.

**Fig. 4.**
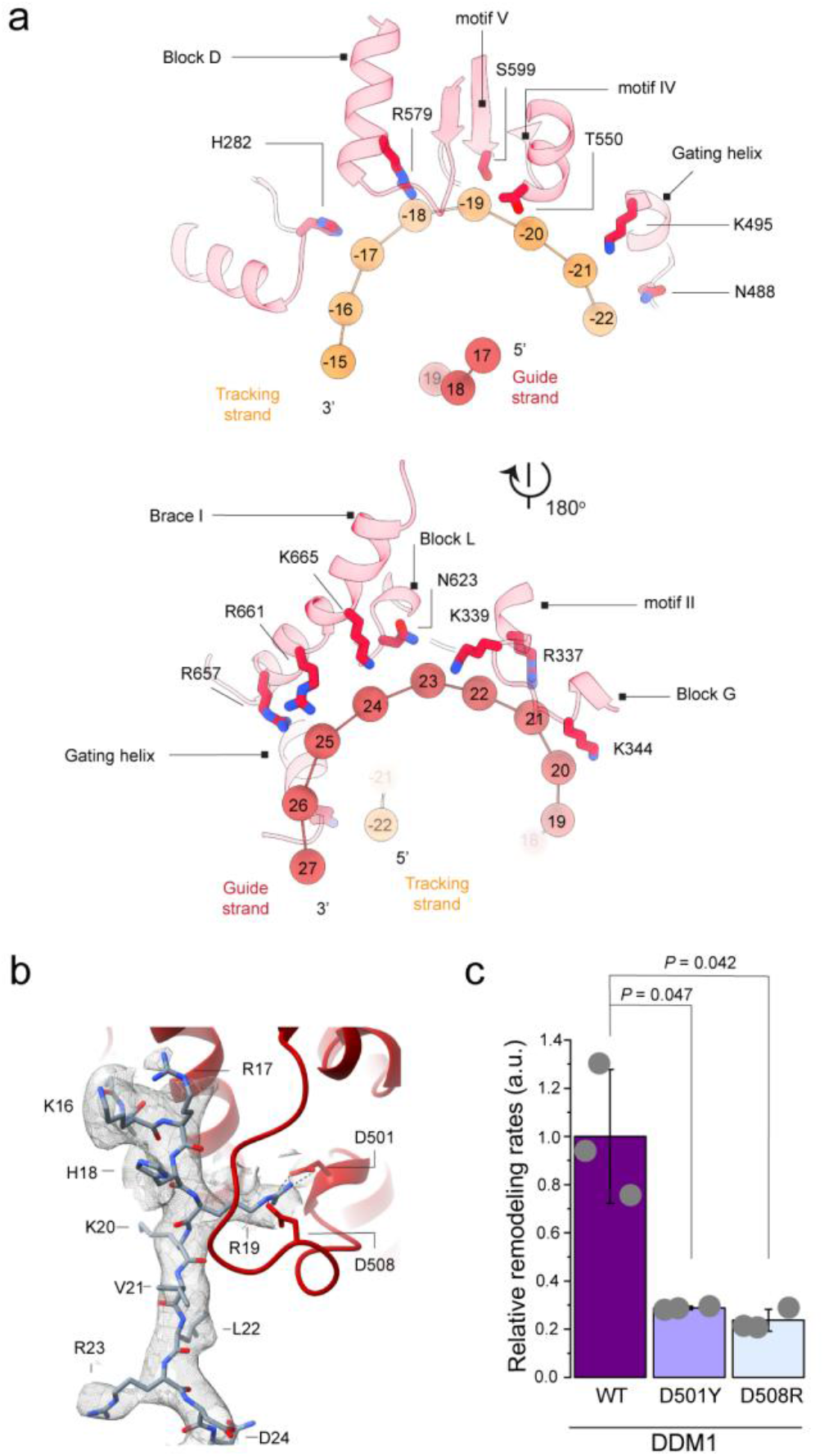
| Interactions of DDM1 motor domain with DNA and histone. **a,** Key residues of DDM1^motor^-DNA interactions are highlighted along with motifs and conserved Snf2 blocks contributing to these interactions. Backbone phosphates are shown as orange spheres. The numbering of the phosphate backbone is relative to the dyad nucleotide **b,** Model-to-map fit of the histone H4 tail interacting with the acidic pocket of ATPase lobe 1. R19 interaction with D501 and D508 of the DDM1 motor domain is highlighted by dashed lines. **c,** Relative rate of remodeling quantified from FRET ratio. Data is mean ± s.d. from three independent experiments. Representative FRET trace is shown in Supplementary Fig. 1b. Statistical comparisons were performed using unpaired student two-tailed t-test. Source data for **c** is provided.

**Fig. 5.**
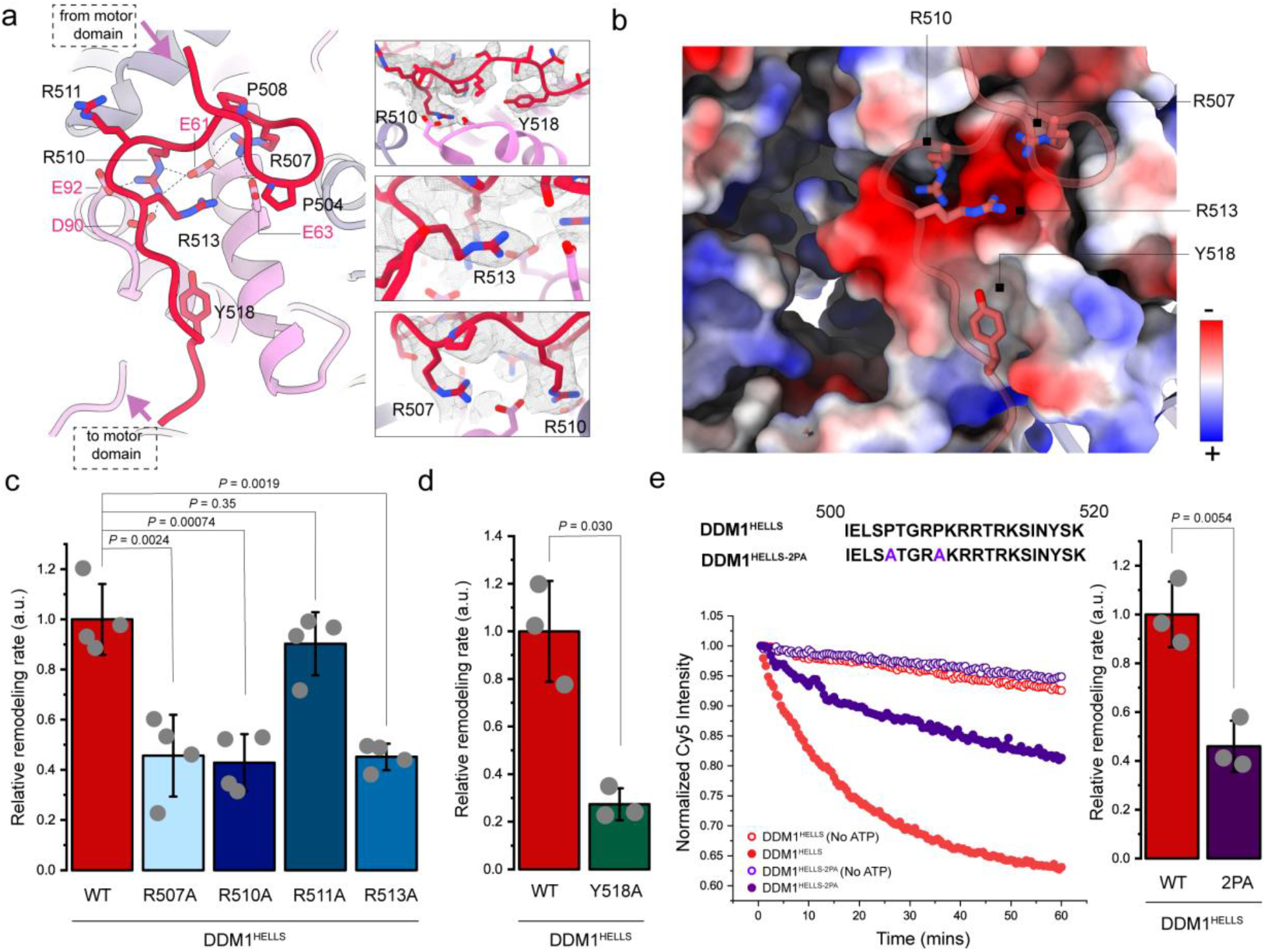
| HELLS midloop interacts with the nucleosome acidic patch. **a,** Structural model of HELLS midloop interaction with the nucleosome acidic patch, showing conserved arginine, tyrosine and proline residues. **b,** Surface electrostatic potential of the nucleosome is depicted. Only key residues involved in acidic patch binding are shown along with Y518 bound to a hydrophobic pocket are shown. **c,** Relative rate of remodeling quantified from FRET traces from four independent measurements of DDM1^HELLS^ wild-type compared to midloop mutants. Data is mean ± s.d. **d,** Quantification of FRET traces from three independent measurements of DDM1^HELLS^ wild-type compared to Y518A mutant. The error bars represent mean ± s.d. **e**, Representative FRET traces comparing DDM1^HELLS^ and DDM1^HELLS- 2PA^ using 0.03 µM nucleosome substrate in the presence or absence of 2 mM ATP (*left*). Quantification of FRET traces after double (DDM1^HELLS^) or single (DDM1^HELLS-2PA^) exponential fit (*right*). Data is mean ± s.d from three independent experiments. All statistical comparisons were performed with unpaired *student’s* two-tailed t-test. Source data for **c**, **d** and **e** are provided.

**Table 1.** Cryo-EM data collection, refinement, and validation statistics.

|  | Nucleosome-bound<br>DDM1-HELLS<br>chimera<br>(EMDB-40569)<br>(PDB 8SKZ) | DDM1 motor<br>(focused)<br>(EMDB-40562) | HELLS midloop-nucleosome complex<br>(focused)<br>(EMDB-40563) |
| --- | --- | --- | --- |
| **Data collection and processing** |  |  |  |
| Magnification | 165,000 | 165,000 | 165,000 |
| Voltage (kV) | 300 | 300 | 300 |
| Electron exposure (e-/Å<sup>2</sup>) | 50 | 50 | 50 |
| Defocus range (μm) | -1.0 to -2.5 | -1.0 to -2.5 | -1.0 to -2.5 |
| Pixel size (Å) | 0.77 | 0.77 | 0.77 |
| Symmetry imposed | C1 | C1 | C1 |
| Initial particle images (no.) | 8507772 | 8507772 | 8507772 |
| Final particle images (no.) | 84067 | 195090 | 195090 |
| Map resolution (Å) | 3.5 | 3.7 | 3.5 |
| FSC threshold | 0.143 | 0.143 | 0.143 |
| Map resolution range (Å) |  |  |  |
| **Refinement** |  |  |  |
| Initial model used (PDB code) | 7ENN, Alphafold2.0 |  |  |
| Model resolution (Å) | 3.5 |  |  |
| FSC threshold | 0.143 |  |  |
| Map sharpening *B* factor (Å<sup>2</sup>) | -50 |  |  |
| Model composition |  |  |  |
| Non-hydrogen atoms | 31859 |  |  |
| Protein residues | 1340 |  |  |
| Nucleotides | 316 |  |  |
| Ligands | 2 |  |  |
| *B* factors (Å<sup>2</sup>) |  |  |  |
| Protein | 131.82 |  |  |
| Nucleotide | 150.88 |  |  |
| Ligand | 143.03 |  |  |
| R.m.s. deviations |  |  |  |
| Bond lengths (Å) | 0.005 |  |  |
| Bond angles (°) | 0.609 |  |  |
| CaBLAM outliers (%) | 0.85 |  |  |
| Cβ outliers (%) | 0.00 |  |  |
| **Validation** |  |  |  |
| MolProbity score | 1.24 |  |  |
| Clashscore | 2.96 |  |  |
| Poor rotamers (%) | 0.61 |  |  |
| Ramachandran plot |  |  |  |
| Favored (%) | 97.04 |  |  |
| Allowed (%) | 2.96 |  |  |
| Disallowed (%) | 0.00 |  |  |

Our structure reveals that DDM1^HELLS^ makes three main contacts with the nucleosome through interactions with the nucleosomal DNA, histone H4 N-terminal tail, and the H2A/H2B acidic patch. The DDM1 motor domain (DDM1^motor^) binds to the superhelical ±2 (SHL2) position of the nucleosome, which is consistent with previously solved chromatin remodeling complexes^51–54^.

### DDM1^motor^-nucleosome interactions

Similar to previously solved SNF2 structures, using ADP.BeFx induces a closed state of the two ATP lobes on the DNA, mimicking the transition state of the ATP hydrolysis cycle. The DDM1^motor^ lobes in this closed state make extensive contacts with the DNA phosphate backbone (Fig. 4a). Mutating three conserved DDM1 residues that are involved in DNA contacts (K339, N488 and T550) to glutamate results in defective nucleosome-simulated ATPase activity (Supplementary Fig. 1a).

A common feature of SNF2 family remodeler–nucleosome binding is the interaction of the H4 N-terminal tail with an acidic cavity of the ATPase lobe 2^48,51,55–57^; this interaction is essential for chromatin remodeling^58^. In our structure, we obtain clear density of the H4 tail interaction with DDM1^motor^ lobe 2, allowing us to unambiguously assign residues G14-R23 (GAKRHRKVLR) (Fig. 4b, Supplementary Video 2). This assignment enables us to determine the exact interaction of critical residues in the H4 tails with the acidic DDM1^motor^ surface and compare the binding mode with other remodelers. Extending from the nucleosome dish-like surface, the H4 tail contacts the major insertion region of ATPase lobe 2 via R19 (Fig. 4b). The major insertion region is a polypeptide between blocks C and K of the SNF2 family enzymes^50^. In most remodelers (except the INO80 family) this insertion is modest and contributes significantly to the H4- interacting acidic cavity of lobe 2. Histone H4 R19 forms bifurcated salt bridges with two DDM1^HELLS^ aspartate residues, D501 and D508, in the major insertion loop (Fig. 4b). To our knowledge, this mode of interaction is unique to DDM1 and is essential for the enzyme’s activity because D501Y or D508R mutations severely impairs the ability of DDM1 to remodel end-positioned nucleosomes (Fig. 4c, Supplementary Fig. 1b). Comparing different ADP.BeFx-bound remodeler structures from distinct families reveals diversity in the mode of interactions with the H4 basic patch, K16-K20 (KRHRK), with ATPase lobe 2. Whereas DDM1 and SNF2H interact mainly via R19 with the insertion loop, ALC1 and CHD4 interact primarily via R17 with acidic residues on a helix between motif IV and block D (Supplementary Fig. 1c).

### HELLS midloop interacts with the nucleosome acidic patch

We observe significant density near the entry-side acidic patch of the nucleosome (Fig. 3a). The HELLS midloop contains a stretch of conserved positively charged residues and two prolines between S503 and S515. We hypothesized that these prolines (P504 and P508) are responsible for the sharp turns in the density near the acidic patch. To test if this density is from the HELLS midloop, we generated a SUMO-tagged protein containing the HELLS midloop polypeptide and conducted direct interaction studies using AlphaScreen. Using a biotin-tagged nucleosome (bio-10N0), we find that the midloop polypeptide directly interacts with the nucleosome (Extended Data Fig. 6a). Importantly, we were able to outcompete this interaction using a synthesized LANA peptide (Extended Data Fig. 6b), which binds the nucleosome acidic patch with high affinity and specificity^59^. Furthermore, in our FRET assays the presence of LANA peptide abolishes the ∼3.5-fold stimulation of nucleosome remodeling seen for DDM1^HELLS^ versus DDM1 (Extended Data Fig. 6c-d). Collectively, these results show that the HELLS midloop stimulates chromatin remodeling through nucleosome acidic patch interactions.

**Fig. 6.**
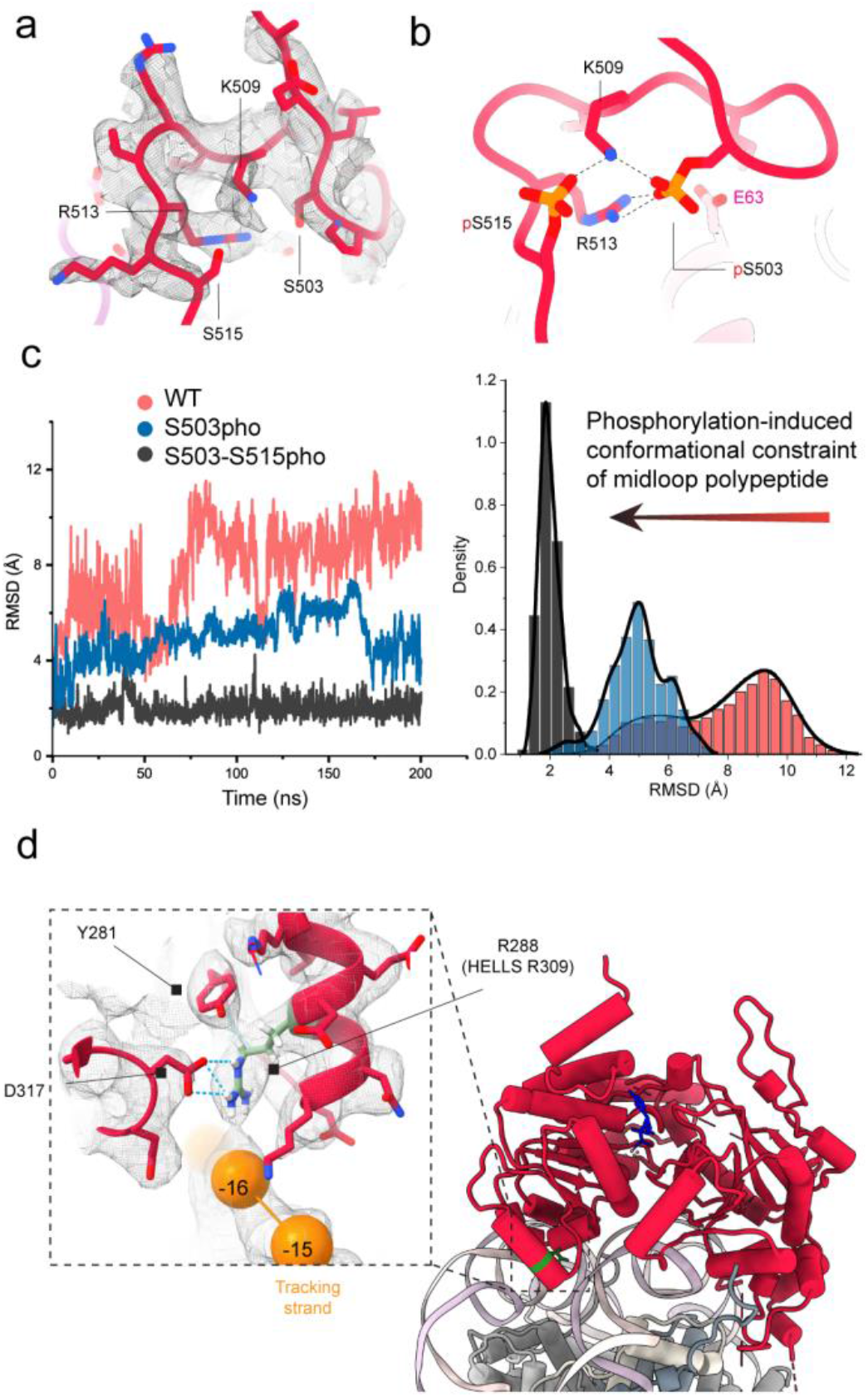
| Post-translational modifications modulate HELLS midloop dynamics. **a,** Model-to-map fit of HELLS midloop, highlighting the S503 and S515 phosphorylation sites. **b,** Model of HELLS midloop following *in silico* phosphorylation of S503 and S515, followed by minimization with ISOLDE. Key residues involved in electrostatic interactions are shown. **c,** Representative RMSD plots of three independently run (Extended Data Fig. 8b) MD simulations comparing wild-type HELLS midloop and phosphorylated models over 200 ns (*left*). Density plot distribution of RMSD fluctuations compared to starting structure (*right*). Distributions were fitted using the Kernel Smooth approach implemented in Origin 2022. **d,** The position of R288 (HELLS R309), methylated in HELLS, is highlighted in green in the DDM1^HELLS^-nucleosome complex. Model-to-map fit of the boxed region to highlight key surrounding interactions (*inset*).

We next modelled a stretch of 20 amino acids of the HELLS midloop (I500 to K520) into the density near the acidic patch, revealing a loop that makes extensive contacts with the acidic patch including interactions through three conserved arginines (Fig. 5a-b, Supplementary Video 3). R510 interacts with the acidic triad of H2A, E92, E61, and D90, forming what is known as the ‘arginine anchor’. R507 serves as a “variant arginine, type I” and interacts with H2A via E61. These two arginine residues are observed commonly in acidic patch-interacting proteins^60^ (Fig. 4b; upper panel). In addition, a third arginine, R513, serves as the rarely observed “variant arginine, type II”, in proximity to H2A E64. A fourth arginine, R511, present on this modelled midloop region does not contact the histones. To confirm the effect of the arginine mutations on nucleosome remodeling, we introduced corresponding mutations into DDM1^HELLS^. Using our FRET-based assay we find that DDM1^HELLS^ with the R507A, R510A, and R513A mutations significantly reduce chromatin remodeling compared to wild-type DDM1^HELLS^, whereas the R511A mutation has no significant effect (Fig. 5c).

In addition to the arginine residues, we observe a highly conserved tyrosine, Y518, in the HELLS midloop that binds into a hydrophobic pocket adjacent to the acidic patch formed by H2A residues L65, N68, A69, L85, A86 and N89 (Fig. 5a,b). A conserved tyrosine wedged into this hydrophobic pocket has recently been seen in other SNF2 enzymes^54,61^. With extensive arginine interactions at the acidic patch, it is unclear why nature evolved this conserved tyrosine packing. In yeast CHD1, a fluorescence quenching assay showed that the conserved arginine residues remain the significant contributors to the binding of a "ChEx" fragment to the acidic patch, and the conserved tyrosine contributed to a lesser extent to binding^54^. This ‘tyrosine anchor’ is also conserved in SWI/SNF and was found to be important for chromatin remodeling^61,62^. To examine the contribution of this tyrosine to chromatin remodeling in HELLS, we generated a DDM1^HELLS^ Y518A mutant and using our FRET-based assay find that it significantly decreases nucleosome remodeling (Fig. 5d, Supplementary Fig. 2), suggesting the ‘tyrosine anchor’ is an evolutionarily conserved binding epitope in SNF2-type remodelers.

Because of the unusual configuration of the acidic patch-binding HELLS midloop compared to modes of interaction by other remodelers (Supplementary Fig. 3, Supplementary Video 3), we examined the importance of the proline-induced turn in nucleosome recognition and remodeling. We generated a DDM1^HELLS^ 2PA mutant containing P504A and P508A, and find that it has a considerable reduction in nucleosome remodelling compared to DDM1^HELLS^ (Fig. 5e). These results suggest that conformational constraint is an important mechanism for HELLS midloop recognition of the acidic patch.

To contextualize our finding of this unique insertion and its role in chromatin remodeling, we performed an evolutionary analysis to ascertain its origin (Extended Data Fig. 7). Using the highly conserved Cterm motif as a fiduciary marker, we searched beyond metazoan organisms for HELLS orthologs. The earliest observation of a polypeptide insertion at the HD2 region is after the split between plants and opisthokonts, which include fungi and animals (Extended Data Fig. 7b). Outside of opisthokonts, no midloop has been observed in any HELLS ortholog implying that the midloop insertion occurred after the plants-opisthokont split but before fungi-holozoa split. Among the acidic patch binding features, the tyrosine anchor evolved very early as it is present in both fungi and holozoan. The evolution of arginine interactions and proline constraints occurred predominantly in the holozoan. To evaluate midloop-nucleosome binding beyond the metazoan, we generated a SUMO-tagged protein containing the midloop polypeptide from *Salpingoeca rosetta*, a choanoflagellate used to study premetazoan evolution^63^, and performed a direct interaction with the nucleosome (Extended Data Fig. 7c-d). The peptide robustly binds the nucleosome pointing to an early origin of the acidic-patch binding property of the HELLS midloop.

**Fig. 7.**
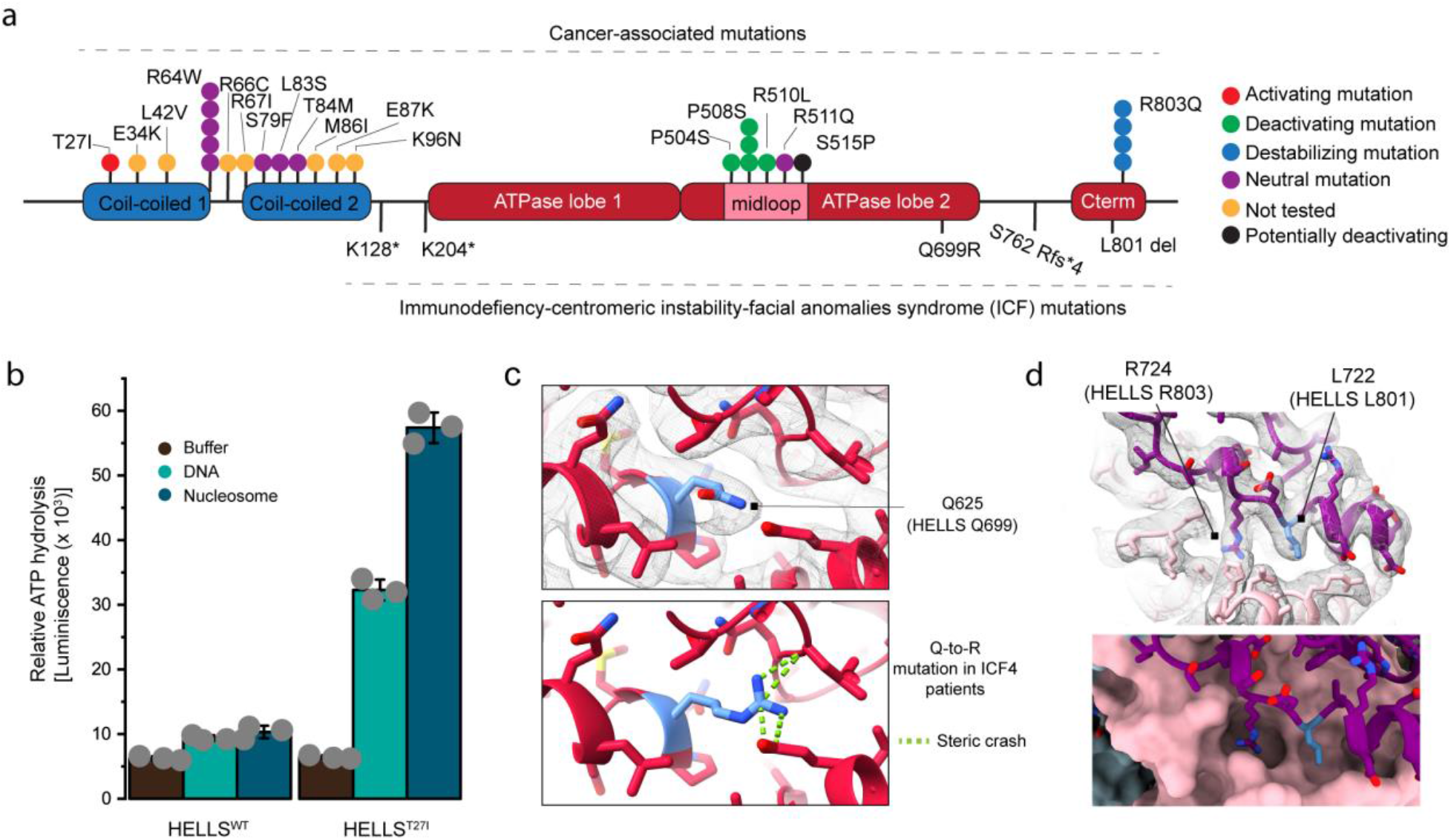
| Disease-associated mutations dysregulate HELLS activity. **a,** *Top,* Patient-derived mutations from cBioportal and COSMIC database are mapped onto regions of interest of the HELLS protein. *Bottom,* ICF4 mutations are mapped onto domains of HELLS. Domains are not drawn to scale. **b,** Endpoint ATP hydrolysis measurements comparing HELLS wild-type and T27I, performed as in Fig 1b. **c,** *Top*, Model-to-map fit of Q625 (HELLS Q699) and *bottom*, model showing the effect of the ICF4 Q-to-R mutation. **d,** Zoomed view showing the relative positioning of the L801 and R803 Cterm residues (purple with the L801 mutant highlighted in blue) bound to ATPase lobe 2 (pink). *Top*, DDM1^HELLS^ model fit into cryo-EM density. *Bottom*, HELLS Cterm shown as cartoon and ATPase lobe 2 shown in surface representation.

### Effect of post-translational modifications on the activity of HELLS

The effect of histone post-translational modifications on the interaction between chromatin and chromatin regulators is a well-studied area of biology and forms the bedrock of modern epigenetics. However, very few studies have mechanistically investigated the effect of chromatin remodeler post translational modifications on their activities. A recent study in cancer stem cells found that HELLS S503 phosphorylation by MAPK1 promotes expression of genes to maintain the cancer stem cell state and drive tumor growth^41^. In addition to S503, S515 is also frequently phosphorylated *in vivo* (Extended Data Fig. 8a) and both residues are observed in the acidic patch-binding midloop of HELLS (Fig. 6a, b). An all-atom molecular dynamics simulation was performed on the midloop structure to investigate the structural effects of phosphorylation. Analysis of the C_α_ atom’s root-mean-square deviation from the initial starting structure reveals a relatively high degree of conformational flexibility of the wild-type HELLS midloop (Fig. 6c, Extended Data Fig. 8b). Phosphorylation of S503, S515 and their combination increased the midloop’s rigidity due to the introduction of salt bridges between the phosphate groups of S503p and S515p to R513 and K509, respectively. In essence, these phosphorylation events preorganize the midloop polypeptide into a similar conformation observed in the bound state (Fig. 5c, Supplementary Video 4). The potential enhancement in binding affinity associated with this preorganization may contribute to the role of HELLS S503 phosphorylation in maintaining cancer stemness^41^.

Recently, HELLS was also shown to be monomethylated at R309 *in vivo* by PRMT5, with a R309A methylation-deficient mutant preventing differentiation and promoting growth of lung cancer stem cells compared to wild-type controls^41^. HELLS R309 is a highly conserved residue in ATPase lobe 1, and analysis of the equivalent DDM1 R288 in our structure revealed that this residue contacts the phosphate backbone of nucleosomal DNA (Fig. 6d). The arginine is constrained by surrounding interactions, leading to the exposure of two terminal nitrogens to solvents. This arrangement also explains why the terminal nitrogen cannot accommodate two methyl groups (asymmetric dimethylation), as one of each of the terminal hydrogens is either sterically occluded or engaged in a salt bridge. *In silico* placement of a methyl group on the terminal nitrogen creates a steric clash with the backbone DNA, which can potentially lead to deficiency in nucleosome binding or remodeling (Supplementary Fig. 3).

### N- and C-terminal regulatory regions of DDM1

The N-terminal region of DDM1, including the coiled-coil, was not resolved in our structure (Extended data Fig. 9a). However, at a lower map threshold, additional densities were observed packed onto the ATPase lobe 1, likely via hydrophobic interactions (Extended Data Fig. 9b-c). These densities can accommodate 1 or 2 helices from the five predicted helices at the unresolved region. The arrangement of these helices is reminiscent of the packing of AutoN against suppH^64^ and postHSA against suppH^48^ in ISWI and yeast Snf2, respectively (Supplementary Fig. 4a). In both cases, these interactions were reported to regulate the enzyme’s activity. To ascertain whether the N-terminus of DDM1 has cryptic regulatory functions like HELLS, we generated DDM1 N-terminus-deleted constructs and compared their activity to wild-type protein (Extended Data Fig. 9d). The removal of the DDM1 N-terminus led to an ∼60% increase in remodeling activity, which is much lower than the ∼3.5-fold stimulation seen for ΔCC1-HELLS.

Comparison between our structure and different SNF2 motor structures bound to ADP.BeFx reveal global differences unique to DDM1^HELLS^ (Supplementary Fig. 4). We observe a protrusion formed by the C-terminus of the DDM1 motor in our structure (Extended Data Fig. 9e, Supplementary Fig. 4b). The Cterm motif present among the DDM1 and HELLS family of chromatin remodelers contains a highly conserved arginine, DDM1 R724 (HELLS R803), which makes extensive contacts to DDM1 ATPase lobe 2 through hydrogen bonds with D613 and T614, and a cation-π interaction with H541 (Extended Data Fig. 9f). These contacts stabilize the C-terminus interaction with the motor protein as a DDM1^HELLS^ R724Q mutation destabilizes the protein leading to substantial aggregation seen during size-exclusion chromatography (Extended Data Fig. 9g). Furthermore, when monomeric fractions of DDM1^HELLS^ R274Q, but not wild-type, is frozen and thawed it either unfolds or aggregates as shown by high initial fluorescence signal in thermal stability assays (Extended Data Fig. 9h).

### Disease-associated mutations in HELLS

HELLS overexpression has been associated with the progression of numerous types of cancers. A search of the CBioPortal and COSMIC databases reveals somatic mutations across the various domains of the protein (Fig. 7a). Here, we focus on mutations that map to the HELLS N-terminal coiled-coil region, the midloop and the C- terminal motif. HELLS T27I maps to the autoinhibitory coiled-coil 1, and we find that similar to our deletion results (Fig 1d) purified HELLS T27I activates ATPase activity in our endpoint assay (Fig. 7b). In some cancers, the two midloop prolines (P504 and P508) are mutated to serine, which based on our results on the proline-to-alanine mutations (Fig. 5e), would likely decrease HELLS nucleosome remodeling. A similar impact on chromatin remodeling is also expected for mutation of the R510 arginine anchor to leucine, which is seen in some cancers. A recurring mutation of the conserved Cterm arginine to glutamine, R803Q, is also seen and, based on our results on the equivalent DDM1 R724Q mutant, we expect it to destabilize the protein (Extended Data Fig. 9g). A structural chromosomal aberration involving HELLS has been observed in several cancers (FusionGDB2 database); for example, in hepatocellular carcinoma, a fusion of CYP2C18 (exon 2) to HELLS (exon 4) replaces the first 145 residues of HELLS with the N-terminus of CYP2C18, eliminating the coiled-coil regulatory elements (Extended Data Fig 10a,b). This fusion is likely to activate HELLS as observed in the ΔCC1-HELLS mutant. However, it must be noted that this fusion also eliminates the HELLS N-terminal nuclear localization signal and further studies are needed to investigate if the fusion protein gets into the nucleus.

Besides the cancer-associated mutations, genetic mutations in HELLS are causative for the disease ICF4, marked by hypomethylation of the genome^11^. Out of the five causative HELLS mutations observed in ICF4 patients (K128*, K204*, Q699R, S762 Rfs*4 and delta-L801) two nonsense mutations, K128* and K204* lead to the elimination of the motor domain (Fig. 7a). The ICF4 HELLS Q699R point mutation is found in a conserved block/motif, L/IV, of SNF2 family remodelers (Extended Data Fig. 2), and computationally modelling this mutation introduces steric collision with neighboring residues that likely destabilize the protein (Fig. 7c). S762 Rfs*4 leads to the elimination of the stabilizing C-terminus. How the in-frame deletion of L801 in the Cterm motif affects the function of HELLS is not known. Analysis of the interaction of the Cterm motif with ATPase lobe 2 reveals that deletion of L801 (DDM1 L722), two residues downstream of R803 (DDM1 L724), has the potential to dislodge the conserved arginine from its pocket leading to destabilization of the protein (Fig 7d).

## Discussion

Orphan chromatin remodelers belong to a subfamily of SNF2 family proteins, which contain diverse and unique regulatory domains not found in the other four major families, include ALC1, CSB/Rhp26, fun30/SMARCAD1, and HELLS/DDM1. ALC1 contains a macrodomain that, in the absence of PAR chains, autoinhibits the ATPase motor^51,65^ and Rhp26 contains a “leucine latch” at the N-terminus that blocks ATP hydrolysis^66,67^. Other chromatin remodelers also use different mechanisms for autoinhibition. In CHD1, an N-terminal chromodomain wedges between the catalytic lobes preventing ATP hydrolysis in the absence of DNA and nucleosomes^44^, and ISWI contains AutoN and NegC motifs that negatively regulate ATP hydrolysis in the absence of substrates^45,46^. Here, we have uncovered a novel autoinhibitory element in the coiled-coil domain of HELLS responsible for the inactivity observed in *in vitro* studies even in the presence of substrates. Even though the coiled-coil region is found in both HELLS and DDM1, only HELLS exhibits strong autoinhibition, with removal of the entire coiled-coil region of DDM1 leading to a relatively slight increase in chromatin remodeling.

The divergence of functions between HELLS and DDM1 is evident from sequence alignment analyses. Residues and secondary structure elements in this region are less conserved between HELLS and DDM1 than in the rest of the protein. Our DDM1^HELLS^ structure reveals the presence of two helices, αH4 and αH5, that are absent in HELLS. These helices are sequestered onto protrusion 1, exposing the coiled-coil domain to solvent. This observation explains why the coiled-coil domain of DDM1 is not observed in our structure and why its deletion does not affect the stability of the protein. This is contrary to HELLS, which we find degrades and aggregates when the entire N-terminus is deleted. Additionally, the fact that the second coiled-coil region of DDM1 has been shown to bind and deposit histone H2A.W further supports the solvent exposure of this region^21^. The N-terminus of HELLS mediates its interaction with numerous binding partners^22,26,37^. Our identification of an N-terminal coiled-coil mediated autoinhibition provides a mechanistic framework for understanding the activation of HELLS by partner proteins^10^.

Similar to other known SNF2-nucleosome complex structures, the interaction of the DDM1 motor domain at the SHL2 is highly conserved. However, in our structure we unambiguously resolve the interaction of the histone H4 N-terminal tail with the DDM1^HELLS^ ATPase lobe 2, allowing us to ascertain the roles of key residues in regulating chromatin interaction. Compared to other remodelers, we observe differences in which H4 arginines (R17 and R19) interact with acidic residues of lobe 2. Post-translational modification of the H4 tail has been shown to regulate chromatin remodeling^68,69^. *In vivo* mutational studies revealed that H4 R17 regulates flowering time in *A. thaliana,* as R17 mutation affects the action of plant ISWI^70^. Differential consequences of R17 and R19 mutations were also observed in *Drosophila* with R17A and R19A affecting H4 K16 acetylation and H3K79 methylation, respectively^71^. We posit that the differences in the observed interaction with the H4 tail could be a means of targeting or blocking the recruitment of remodelers to specific functional niches. However, we cannot exclude the possibility that differences may be due to the poor density observed for this region in other complexes compared to our DDM1^HELLS^-nucleosome structure.

The acidic patch of the nucleosome is a hotspot for protein-protein interaction^60^. SNF2 family remodelers use auxiliary motifs, normally located N- or C-terminal to the ATPase motor, to bind the nucleosome acidic and stimulate chromatin remodeling^72,73^. We have identified a unique insertion in the ATPase lobe 2 of HELLS that has evolved to stimulate remodeling activity via interaction with the acidic patch, with proline-induced conformational constraints used to facilitate binding. Furthermore, our results suggest that phosphorylation of S503 and S515 in the midloop accentuates these conformational constraints by establishing new salt bridges with K509 and R513. This leads to more efficient binding and potentially enhanced chromatin remodeling. Our findings provide a new paradigm for understanding and investigating acidic patch interactions beyond the well-known arginine anchors.

In lung cancer stem cells, phosphorylation of S503 by MAPK1 promotes the expression of cancer stem cell markers to maintain stemness^41^. The effect of this phosphorylation has been shown to crosstalk with R309 methylation, with methylated R309 driving differentiation of the lung cancer stem cells. Our results from molecular dynamics simulation suggest that phosphorylation may enhance HELLS nucleosome remodeling activity, whereas our structure reveals that R309 methylation will hinder efficient interaction of HELLS to the nucleosome SHL2. These observations provide mechanistic insights into the opposing effect of these two modifications on the maintenance of cancer stem cell state. In addition to chromatin remodeling, HELLS has been shown to deposit the macroH2A histone variant into nucleosomes to repress the expression of target genes^18,20^ and promote homologous-recombination dependent restart of stalled replication forks^19^. A recent study with SWR1C highlighted the importance of the nucleosome acidic patch in the exchange of the H2A.Z/H2B dimer^74^. A putative arginine anchor in the Swc5 subunit interacts with the acidic patch to mediate histone exchange. The identification of a HELLS midloop capable of binding the acidic patch may imply a similar role in the deposition of the macroH2A/H2B dimer.

The C-termini of SNF2 and related remodelers contain a regulatory domain, NegC in ISWI, SnAc in SNF2, and a C-terminal bridge in CHD, that negatively impact chromatin remodeling^44,57^. These domains are absent in HELLS/DDM1. Instead, we find that a family-defining motif, which we call Cterm, binds and stabilizes as an “arch” structure onto the ATPase motor lobe 2. Holding together the C-terminus-ATPase lobe 2 interaction is a highly conserved arginine (HELLS R803, DDM1 724), mutation of which we find to destabilize the protein. This observation is similar to findings with *M. thermophila* Snf2, where a SnAc C-terminal domain binds to and stabilizes the ATPase lobe 2^75^. The evolutionary convergence and dependence on this C-terminal arginine for stability make it a hotspot for disease regulation.

Taken together, we took a systematic approach to reveal important epitopes for chromatin remodeling of an elusive family of heterochromatin remodelers that occupy a unique functional niche in the cell. Our study provides mechanistic insight into the function and regulation of the HELLS/DDM1 family of remodelers and a structural framework to rationally develop small molecules to therapeutically target HELLS.

## METHODS

### Cloning and purification of HELLS, DDM1 and SNF2H proteins

A pENTR 223.1 plasmid containing the ORF for Homo sapiens HELLS was obtained from transomic (Clone ID: BC146308) and used as a template for PCR. To amplify the gene encoding for full-length HELLS, forward and reverse primers were designed to allow its incorporation into a pET His6 TEV cloning macrolab vector via ligation independent cloning (LIC, https://qb3.berkeley.edu/facility/qb3-macrolab/). The plasmid was electroporated into Rosetta™ 2 cells (Millipore Sigma) for protein expression. To produce the recombinant N-terminally 6xHis-tagged HELLS, liquid cultures were incubated in ampicillin (100 μg/mL) and chloramphenicol supplemented (25 μg/mL) LB medium at 37°C with shaking at 200 rpm overnight. The culture was transferred into a 2-4 L Terrific Broth medium supplemented with antibiotics. At an optical density (OD) of 0.6 isopropyl (thio)-β-D-galactopyranoside (IPTG, 0.6 mM) and MgCl2 (5 mM) were added. Following overnight incubation at 16 °C, cells were harvested at 7,500 x g for 15 min at 6°C. To purify full-length HELLS, cell pellets were resuspended in lysis buffer (20 mM HEPES, pH 6.8, 750 mM NaCl, 10 % glycerol, 2 mM MgCl2, 2 mM PMSF, 1 mM DTT, 0.1% Triton X-100, 0.2 mg/mL lysozyme, 0.1 mg/mL DNase II, 10 mM imidazole and 1 SIGMAFAST™ Protease Inhibitor Cocktail Tablet (Millipore Sigma) and incubated on ice for 30 mins. Subsequently, the suspension was lysed on ice by sonication for 5 x 45 seconds. Lysates were separated by centrifugation at 15,000 x g for 35 min. The supernatant was filtered (0.45 μm; Millipore) and mixed with 3 mL of Ni-NTA resin (Qiagen) for 1.0 h at 4 °C and then applied to a gravity flow column. The matrix was washed with 50 column volumes (CV) of washing buffer (20 mM HEPES pH 6.8, 750 mM NaCl, 10 % glycerol, 2 mM MgCl2, 2 mM PMSF, 1 mM DTT, 0.05% Triton X-100, 30 mM Imidazole) and the protein eluted with an imidazole-gradient (50-500 mM). Fractions containing full-length HELLS were pooled and Tobacco Etch Virus (TEV) protease (1 part TEV to 100 parts HELLS) added to cleave the N-terminal His-tag during overnight dialysis into a no-imidazole buffer A (20 mM HEPES pH 6.8, 500 mM NaCl, 10% glycerol). The cleaved protein was applied to a 5 mL HisTrap HP pre-packed column (Cytiva). A portion of HELLS flowed-through the column along with contaminating *E. coli* proteins and degradation products. A portion of cleaved HELLS bound to the column and was eluted in a homogeneous fraction in a step-wise procedure using imidazole concentrations, formed by mixing buffer A and buffer B (20 mM HEPES pH 6.8, 500 mM NaCl, 10% glycerol, 500 mM Imidazole), between 5-25 mM. Collected protein was concentrated using Vivaspin protein concentrator spin column (10 kDa MWCO, Cytiva) and subsequently applied to a HiLoad 16/600 Superdex 200 pg preparative SEC column (Cytiva) equilibrated with 20 mM HEPES pH 6.8, 500 mM NaCl, 2 mM DTT, 10% glycerol. Peak fractions were concentrated, flash frozen and stored at - 80°C. Truncation and single amino acid mutations in HELLS were generated by one-step site-directed mutagenesis and protein purified similar to wild-type HELLS.

Arabidopsis thaliana DDM1 was cloned from a pENTR/SD-dTopo vector containing the ORF of the DDM1 gene (Arabidopsis Biological Resource Center, OSU, USA) using LIC. The PCR product was subcloned into a pET His6 TEV LIC cloning vector (2B-T) and protein was overexpressed following the same protocols detailed for HELLS above. Cells were harvested and lysed in a buffer containing 20 mM Tris-HCl pH 7.5, 500 mM NaCl, 10% glycerol, 2 mM PMSF, 1 mM DTT, 0.1% Triton X-100, 0.2 mg/mL lysozyme, 0.1 mg/mL DNase II, 10 mM imidazole and 1 SIGMAFAST™ Protease Inhibitor Cocktail Tablet (Millipore Sigma) and incubated on ice for 30 mins. After sonification, the lysate was centrifuged at 15,000 x g for 35 mins and the soluble fraction incubated with 3 mL of Ni-NTA beads (Qiagen) for 1 hour at 4°C. Bound proteins were eluted with 20 mM Tris-HCl pH 7.5, 500 mM NaCl, 10% glycerol, 2 mM PMSF, 1 mM DTT buffer supplemented with an increasing amount of imidazole. Eutions containing DDM1 were pooled, digested with TEV protease, and dialyzed overnight into a low salt buffer (20 mM Tris-HCl pH 7.5, 150 mM NaCl, 10% glycerol, 2 mM MgCl2). The sample was then diluted and applied to a 5 mL HiTrap Q HP column (Cytiva). The protein was eluted with salt gradient (50-500 mM) and dialyzed into 20 mM HEPES pH 7, 100 mM NaCl, 10% glycerol, 2 mM MgCl2, for 6 hours at 4°C and then applied to a Capto HiRes S column (Cytiva). After a linear salt elution, the DDM1 peak was collected and applied onto a HiLoad 16/600 Superdex S200 pg size exclusion column. The monomeric peak was collected and concentrated with a 50 kDa spin column, flash-frozen and stored at -80°C.

To generate DDM1^HELLS^, Gibson assembly was used to subclone a region corresponding to the HELLS mid loop (I500-E553), between G479 and Q480 of wild-type DDM1. The resulting clone was transformed, expressed, and purified using the wild-type DDM1 protocol. Expression constructs for all DDM1 and DDM1^HELLS^ point, and deletion mutations were created using site-directed mutagenesis and expressed and purified as for wild-type protein.

Full-length human SNF2H was cloned into a pET His6 TEV LIC cloning vector (2B-T) and expressed using the HELLS protocol in a T7 Express lysY/I q Competent E. coli cell (NEB). To purify the expressed protein, cells were sonicated in a 20 mM HEPES pH 7.5, 500 mM NaCl, 10% glycerol, 10 mM Imidazole, buffer supplemented with 0.2 mg/mL lysozyme, 0.1 mg/mL DNase II, 10 mM Imidazole, and 1 SIGMAFAST™ Protease Inhibitor Cocktail Tablet (Millipore Sigma). After centrifugation (15,000 x g for 35 mins), the supernatant was applied to a 5 mL HisTrap column, washed, and eluted with an increasing concentration of Imidazole. The eluted protein was pooled, and the His-tag cleaved with TEV protease. After overnight dialysis, the sample was applied to an 16/600 Superdex S200 column, concentrated and stored at -80 C.

### Purification of histones

A pET29a-YS14 plasmid containing *Xenopus laevis* histones was obtained as a gift from Jung-Hyun Min (Addgene plasmid # 66890). H2A-H2B was subcloned into pET His6 TEV LIC cloning vector (2B-T) using PCR amplification and LIC, and H3-H4 was subcloned into pET His6 TEV LIC cloning vector (2A-T) with a C-terminal His6 tag using a similar protocol. These plasmids were transformed into T7 Express lysY/I q Competent E. coli cells (NEB) and overexpressed. To purify histones, pellets were resuspended in 20 mM Tris/HCl pH 8, 2 M NaCl, 3mM β-mecaptoethanol, 1 mM PMSF supplemented with 1 SIGMAFAST™ Protease Inhibitor Cocktail Tablet (Millipore Sigma). The suspension was sonicated and centrifuged at 15,000 x g. Ni-NTA beads were added to the soluble fraction and the mixture incubated for 1 hour at 4°C. Using gravity flow, the beads were washed with 30 CV of binding buffer supplemented with 20 mM imidazole and the protein eluted with three rounds of 2 CV elution buffer (binding buffer with 300 mM imidazole). The histone dimers and tetramers are further purified with size exclusion chromatography.

### Nucleosome preparation

To form the histone octamer, histone dimers and tetramers are mixed in a ratio of 1.2:1 with TEV protease, followed by overnight dialysis into fresh binding buffer. Size exclusion chromatography was used to separate histone octamer from other histone species. The octamer is concentrated and dialyzed into a 50% glycerol buffer for storage at -20°C. DNA for nucleosome formation was generated using PCR and precipitated with isopropanol. The pellet was redissolved in water and further purified using a Superose 6 size-exclusion chromatography column (Cytvia). The eluted peak is concentrated and equilibrated into nucleosome formation buffer (20 mM Tris/HCl pH 7.5, 2 M NaCl, 1 mM DTT. Histone octamer and DNA were mixed in equimolar amounts and stepwise salt dialysis was used to form the nucleosomes. Nucleosomes were quantified with nanodrop using extinction coefficient calculated from protein and DNA.

### End-point ATPase assays

End-point ATPase assays were performed with the ADP- Glo^TM^ kinase kit^76^. Briefly, 0.2-0.4 μM of protein were incubated with 500 μM Ultra-Pure ATP in 20 mM HEPES pH 7, 50 mM NaCl, 10% glycerol, 5 mM MgCl2, 0.01% Triton-X100 for 1 hour at room temperature in a volume of 5 µL. Proteins were pre-incubated with 0.1-0.2 μM 601 DNA or 30N30 or 0N50 nucleosomes. After incubation, 5 µL ADP-Glo^TM^ Reagent was added and incubated for 40 mins to remove ATP that was not hydrolyzed. The ADP by-product of the ATPase activity is converted to ATP and subsequently converted to light by the addition of 10 µL Kinase Detection Reagent. The luminescence is measured on a BMGlabtech PHERAstar FSX microplate reader.

### NADH-coupled ATPase assay

NADH-coupled assays were performed as previously published^77^. Briefly, 0.2 -0.4 μM of proteins in assay buffer, 20 mM HEPES, pH 7.5, 50 mM NaCl, 10 mM MgCl2, 5 % glycerol, 1 mM DTT was supplemented with 200 μM NADH, 25 units/μL lactate dehydrogenase, 25 units/μL and 2 mM phosphoenolpyruvate. Nucleosome (0.05 μM) and DNA (0.1 μM), when indicated were added to the mixture. To initiate reactions, 2 mM ATP is added and absorbance at 340 nM was monitored on a BMGlabtech PHERAstar FSX microplate reader.

### Nucleosome remodeling using ensemble FRET

To label nucleosome for bulk FRET measurement, K119C H2A was site-specifically conjugated with Cy3 maleimide (Cytiva) as previously described ^47^. Labelled H2A was assembled with purified H2B, H3 and H4 to form octamer^78^. PCR was used to generate an end labelled Cy5 DNA with 50 nucleotides at the exit side. End-positioned nucleosomes (0N50) were reconstituted as described above. To measure kinetics of remodeling, the Cy3 fluorophore is excited at 540 nm and emission measured at 575 nm and 670 nm for Cy3 and Cy5, respectively with a BMGlabtech PHERAstar FSX microplate reader. 20 µL reaction mixtures consisting of 1.0 μM of enzyme, 0.03 μM nucleosome and, when indicated, 100 µM LANA peptide, were mixed in 20 mM HEPES pH 7.0, 50 mM KCl, 10% glycerol, 10% glucose, 5 mM MgCl2, 0.01% TritonX-100. Remodeling was initiated by the addition of 1 mM ATP at 30 °C with an onboard injector. Kinetic measurements were analysed with Origin software.

### Gel-based nucleosome remodeling assay

Restriction access accessibility assays were performed under single turnover conditions as previously described^79,73^. Proteins (0.4 μM) were mixed with nucleosome (50N66, 0.03 μM - Epicypher) in 20 mM Tris-HCl pH 7.5, 50 mM KCl, 10% glycerol, 0.1 mg/mL BSA, 10 mM MgCl2 and 0.01% TritonX-1. 100. The reaction was incubated at 30°C for 30 mins in the presence of 2 units of DpnII. Remodeling was initiated by the addition of 2 mM ATP and indicated timepoints were taken and quenched with stop buffer (10 mM Tris-HCl pH 7.5, 40 mM EDTA, 10% glycerol, 0.6% SDS, 50 μg/mL proteinase K). The samples were deproteinated by incubation at 55°C for 30 mins. Fractions were resolved by native PAGE (5% TBE, 0.5 X), stained with GelRed^TM^ and visualised with a Biorad ChemiDoc Imaging system.

### Fluorescence polarization DNA binding assay

DNA binding activity of DDM1 wild-type and mutants were quantified with end-point fluorescence polarization assays. A 5’- FAM-labelled 50-mer DNA (Integrated DNA Technologies) was used as a substrate at 5 nM. Increasing concentration of proteins from 0 to 3000 nM were incubated with fixed concentration of substrate in 20 mM HEPES pH 7.5, 50 mM KCl, 10% glycerol and 1 mM DTT for 30 min at room temperature. Polarization was measured with a BMGlabtech PHERAstar FSX microplate reader. Curves were fitted with a quadratic binding equation as implemented in Origin 2022 software.

### Alphascreen assay

Alphascreen assay was performed in the Greiner 384-well small-volume microplate. AlphaScreen Histidine detection kit (PerkinElmer) consisting of streptavidin donor beads and nickle chelate (Ni-NTA) acceptor beads were used at final beads concentration of 20 μg/ml. Biotinylated nucleosome (20 nM) and 6Xhis-tagged protein (varied concentration) were mixed and incubated for 30 mins at room temperature in 10 μl of binding buffer (20 mM HEPES, 120 mM NaCl, 10 % glycerol, 0.1 % Triton X- 100, 0.5% BSA) followed by addition of 10 μl donor and acceptor beads for conjugation. The mixture was further incubated for 30 mins at room temperature and AlphaScreen measured on the BMGlabtech PHERAstar FSX microplate reader. Donor beads were excited at 680 nm and the subsequent emission read at 615 nm.

### Mass photometry

Precision Cover glasses (ThorLabs) were cleaned with MilliQ water and 100% isopropanol and dried with compressed air and placed in 3 mm x 1 mm plastic gaskets (Grace Bio-labs). For calibration, 15 μL of PBS buffer was loaded into designated wells in the gaskets and, after autofocusing, 2 μL of BSA was mixed to a final concentration of 50 nM. A one-minute video was acquired and processed with AcquireMP and DiscoverMP in the Refeyn Two MP system. For DDM1 and DDM1^HELLS^, 2 μL of individual proteins were diluted into 15 µL of PBS solution to a final concentration of ∼40 nM. Data was collected similarly to the BSA standard. The amount of particle impact events were analyzed using ratiometric contrast, and molecular weights were estimated based on BSA calibration curves.

### Phylogenetic analysis

Sequences for analysis were fetched via blast from GenBank to broadly represent members of the metazoa. Alignment was performed with CLUSTER Omega and MUSCLE as implemented in Jalview v.2^80^. Aligned sequences were manually trimmed. Sequences were imported into MEGA11^81^ and Maximum Likelihood method and JTT matrix-based models^82^ were used to build the evolutionary sequence history of HELLS. Initial tree(s) for the heuristic search were obtained automatically by applying Neighbor-Join and BioNJ algorithms to a matrix of pairwise distances estimated using the JTT model, and then selecting the topology with superior log likelihood value. The reliability of the selected topology was tested by performing 500 bootstrap replicates^83^.

### Molecular dynamics

To stimulate the dynamics of the HELLS midloop, the initial structure was extracted from the experimentally determined complex reported in this work. In silico phosphorylation of S503 and S515 using the SP2 patch and subsequent preparation of structures for molecular dynamics were performed in CHARMM-GUI^84^ using the CHARMM36m forcefield^85^. To perform the stimulation, the peptides were placed in a cubic box with a dimension of 49 Å x 49 Å x 49 Å and solvated with TIP3P water and 150 mM NaCl. Periodic boundary conditions were applied, and with Particle Mesh Ewalds electrostatics. The system was minimized using the method of steepest decent. A temperature of 303.15 K was maintained with the Nose-Hoover coupling method with a tau-t of 1 ps, and the Parrinello-Rahman algorithm was used to maintain the pressure coupling. Hydrogen bonds were constrained using LINCS^86^. Three independent 200 ns production runs were performed in GROMACS 2022.1^87^ with 2 fs time steps. RMSD analysis was performed in GROMACS 2022.1.

### Thermal shift assay

Wild type DDM1^HELLS^ and DDM1^HELLS^ R724Q were incubated with SYPRO Orange (2X) at final protein concentration of 5 µM. PCR protocol was executed per published protocol^88^.

### Cryo-EM sample preparation and data collection

To form the complex between DDM1^HELLS^ and nucleosome (30N15), the individual species were mixed in a buffer containing 20 mM HEPES pH 7, 50 mM NaCl, 2 mM MgCl2 and 3 mM ADP.BeF3 at final concentrations of 3 µM (nucleosome) and 30 µM (DDM1^HELLS^). The complex was allowed to equilibrate at room temperature for 30 min after which it was crosslinked with 0.05% glutaraldehyde on ice for 5 mins. The crosslinking was quenched by the addition of 125 mM Tris-HCl. After centrifugation, 3 µl of a 1:1 dilution sample was blotted on a glow-discharged (30 s in air) C-flat^TM^ Holey Carbon grid for 1 s with a blot force of -5 at 8°C and 100% humidity using the FEI Vitrobot Mark IV. The sample was rapidly plunge-freeze in liquid ethane and stored for screening. After screening and preliminary data collection on a 200 kV Glacios, a 24-hour data collection was performed on 300kV FEI Titan Krios equipped with Selectris energy filter with Falcon IV camera. Data were collected at a magnification of 81,000 x at a physical pixel size of 0.77 Å with a defocus range of -2 µm to -1 µm. The total electron dose was 50 e/A2, and 128 internal frames for 896 EER fractions.

### Cryo-EM image processing

Electron events recorded in the EER files were converted into stack of images without up-sampling in cryoSPARC 3.3.2^89^. A total of 7,808 movies were motion-corrected and dose-weighted using patch motion correction. The contrast transfer function was estimated with patch based CTF estimation. The resulting micrographs were manually curated and subjected to automatic particle picking with blob picker followed by a template-based picking in cryoSPARC. After removal of duplicates, 850,7772 particles were extracted with box of 448 pixels and binned fourfold. The resulting particles were subjected to iterative 2D classification, Ab-Initio reconstruction and heterogenous refinement. A subset of 417,256 refined particles were re-extracted at 448 pixels and further refined using non-uniform refinement. A reference-free 3D classification followed by heterogeneous was performed to further separate out the “good” particles. A second-round non-uniform refinement was performed to generate a consensus map at 3.54 Å. At a low threshold, the consensus map revealed an additional density in the motor domain, suggesting that it is a mixture of singly-and doubly-bound complexes. A second round of reference-free 3D classification was performed to separate the two complexes. The singly-bound map was subjected to a third round of non-uniform refinement followed by a local and global CTF refinement. This was followed by a fourth non-uniform refinement and a local refinement to yield a final reconstruction at 3.53 Å. To improve the resolution of regions of interest and to aid in model building, the consensus map was subjected to two separate focus refinement. A soft mask covering the motor domain was applied and region was subtracted from the consensus particle stack followed by local refinement. The density in the vicinity of the acidic patch was improved also improved with similar approach. All sharpening, when needed, was performed within cryoSPARC.

### Model building

To build the atomic model of DDM1^HELLS^-nucleosome complex, the nucleosome was extracted from an existing structure (PDB ID 7ENN). The initial model for DDM1^HELLS^ was generated using Alphafold 2.0^90^ as implemented in Google Colab. Both nucleosome and DDM1^HELLS^ models were rigid body fitted into the singly-bound density map in UCSF chimeraX^91^. The resulting complex was MDFF fitted using ISOLDE^92^ followed by manual adjustments in Coot^93^. The model was subjected to real-space refinement in PHENIX^94^. Iterative runs of ISOLDE, Coot and PHENIX were run using combination of maps from focused refinement and singly-bound reconstruction to further refine the geometry of the model. The quality of the model was validated with Molprobity^95^. Final coordinates have been deposited into RCSB PDB (8SKZ).

## DATA AVAILABILITY, SUPPLEMENTARY MATERIAL STATEMENT

The atomic coordinates of the DDM1^HELLS^-nucleosome complex have been deposited to the RCSB Protein Data Bank with PDB ID 8SKZ. The cryo-EM Coulomb potential maps were deposited in the Electron Microscopy Data Bank as EMD-40569 (DDM1^HELLS^- nucleosome complex), EMD-40562 (focused map of DDM1^motor^ bound to nucleosome), EMD-40563 (focused map of HELLS midloop bound to nucleosome).

## CONTRIBUTIONS

WN designed and performed all experiments. All figures and images in this study were generated by the authors using indicated drawing software and are not subject to copyright elsewhere. WN is co-supervised by GJW and AAG. All authors contributed to data analysis and manuscript preparation.

## EXTENDED DATA FIGURES

**Extended Data Fig. 1.**
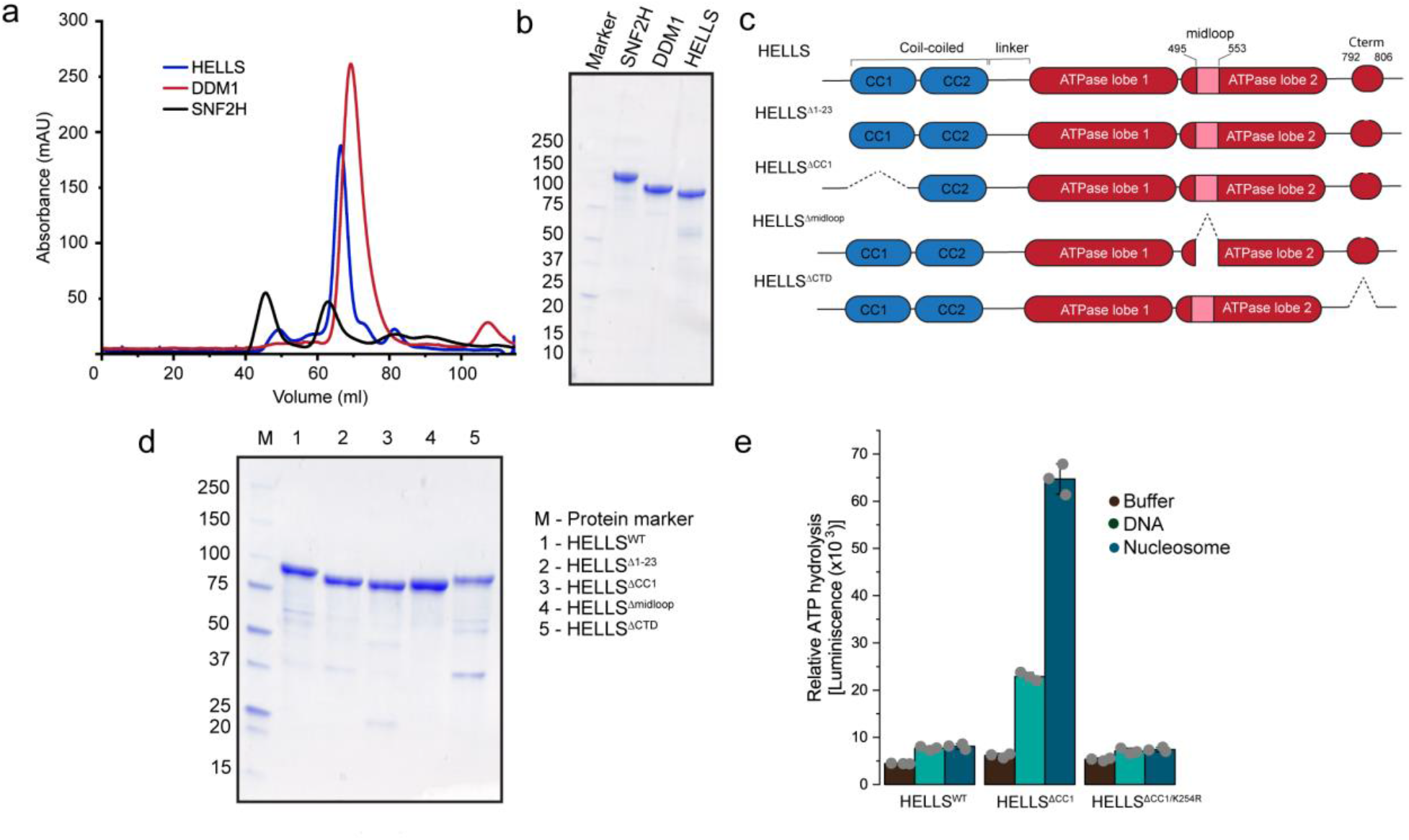
| Identification of a HELLS autoinhibition domain. **a,** Size-exclusion chromatography of wild-type HELLS, DDM1 and SNF2H proteins using Hiload 16/600 Superdex^TM^ 200 pg column. **b,** Coomassie-stained SDS-PAGE of purified proteins. **c,** Scheme for sequential deletion of the internal HELLS motifs. **d**, Coomassie-stained SDS-PAGE of purified mutant proteins. **e,** Relative ATPase activity of wild-type HELLS, HELLS^ΔCC1^ and HELLS^ΔCC1^ K254R using ADP-Glo assay. Data are mean ± s.d. (n=3 technical replicates)

**Extended Data Fig. 2.**
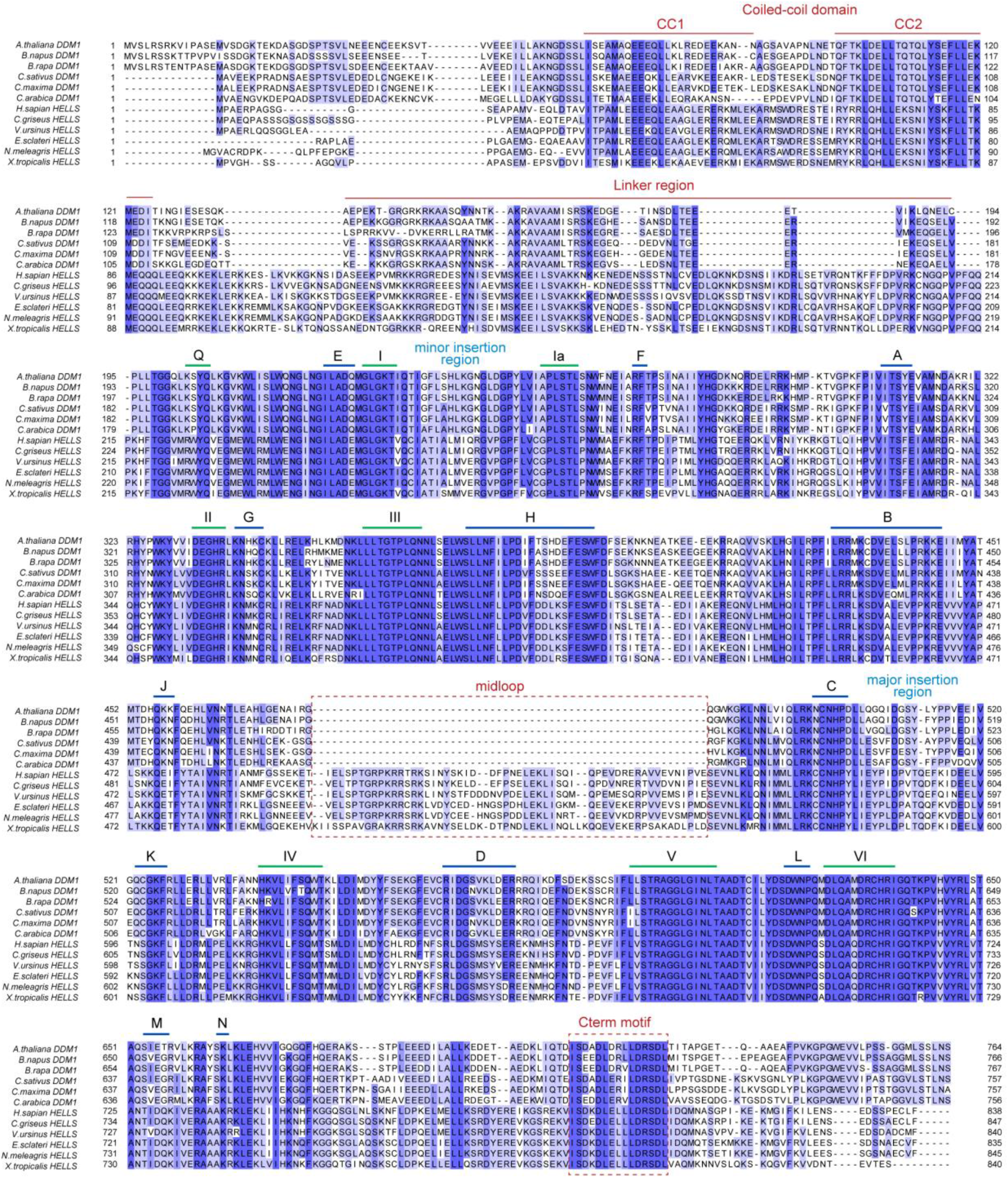
| Sequence alignment of DDM1 and HELLS homolog proteins. Unique features of DDM1/HELLS proteins absent in other Snf2 chromatin remodellers are highlighted with dark red lines and boxes along with labels. Green lines and corresponding label highlight conserved helicase motifs. Blue lines and corresponding label highlight other conserved blocks of Snf2 remodellers.

**Extended Data Fig. 3.**
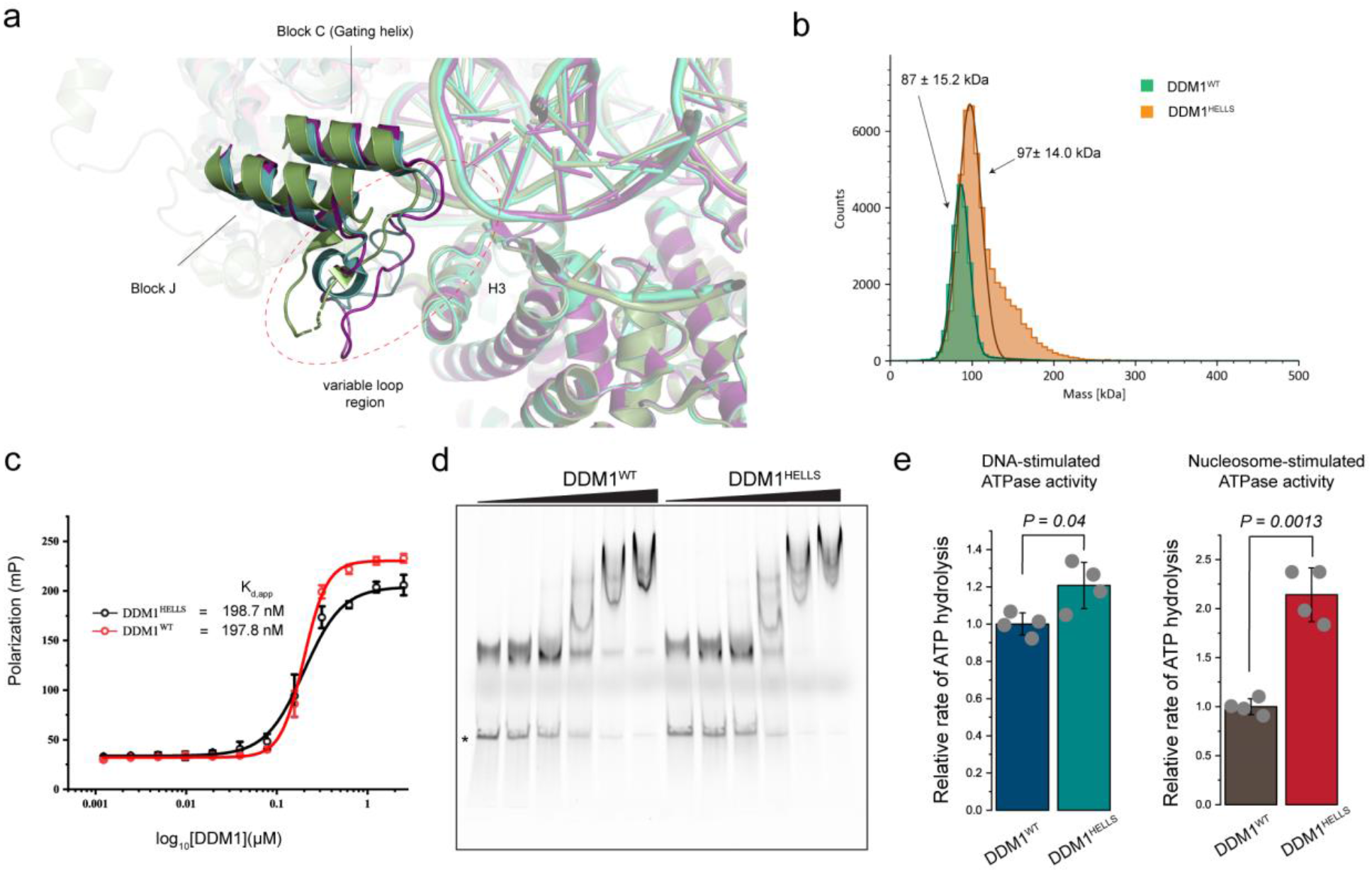
| Generation of DDM1-HELLS Chimera. **a,** Structural and sequence alignment of Snf2 enzymes highlighting protrusion 2 of the ATPase lobe 2. BRG1 (PDB ID 7VDT, Smudge), SNF2h (PDB ID 6NE3, deep purple) and CHD4 (PDB ID 6RYR, light teal) are aligned. **b,** Mass distribution histogram for DDM1 and DDM1^HELLS^. **c,** Fluorescent polarization saturation DNA binding curve for DDM1 and DDM1^HELLS^. FAM-labelled 50-mer DNA (5 nM) was incubated with increasing concentrations of protein for 30 mins at 25^°^C**. d,** Increasing concentration of DDM1 proteins was titrated against fixed concentration of Cy5-labelled DNA (10 nM). **e,** NADH-coupled ATP hydrolysis assay comparing DDM1 and DDM1^HELLS^ activity in the presence of DNA and nucleosome. Data are mean ± s.d. of four technical replicates). Statistical comparisons were performed with unpaired student’s two-tailed t- test. Source data for **c** and **e** are provided.

**Extended Data Fig. 4.**
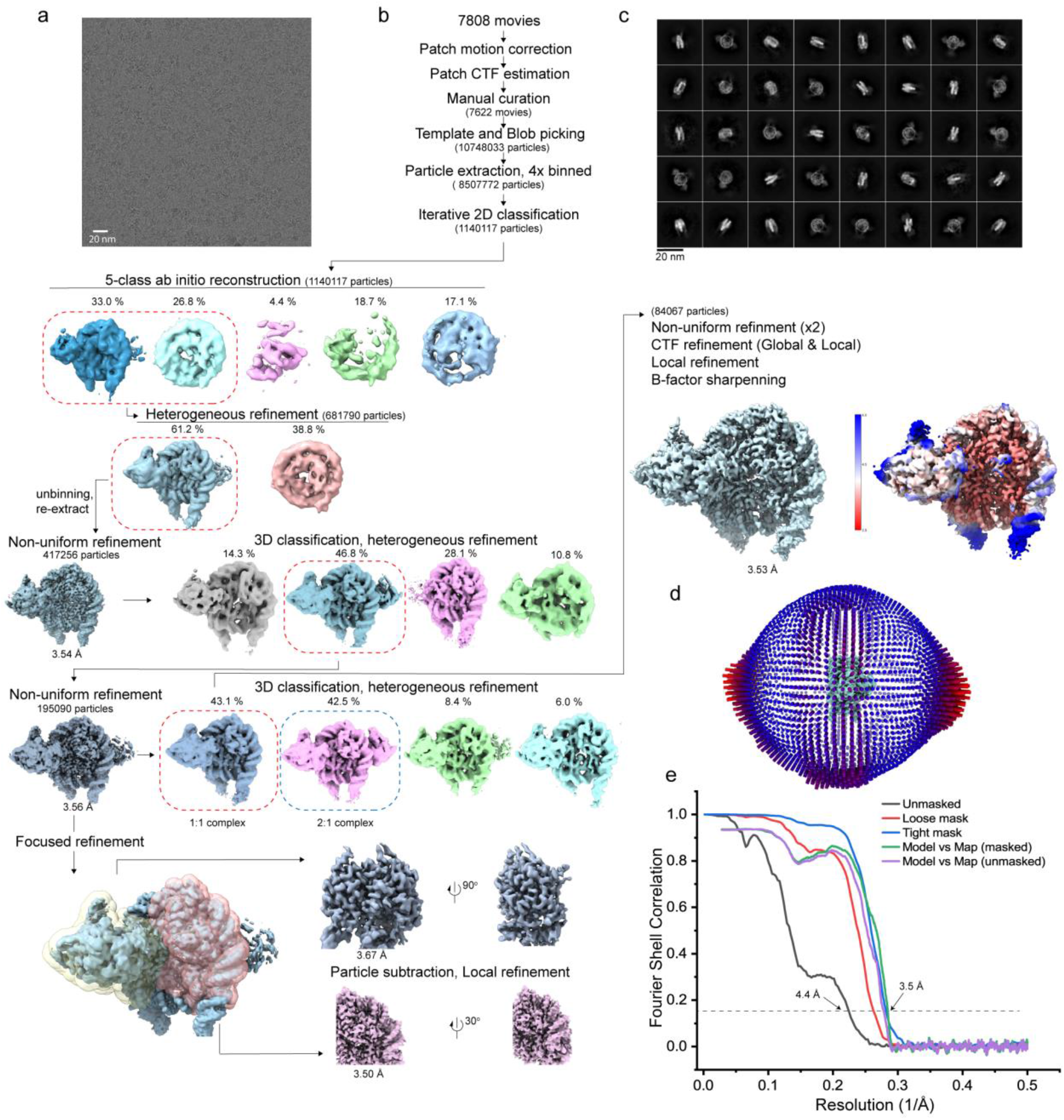
| Cryo-EM analysis of the DDM1^HELLS^-nucleosome complex. **a,** A representative micrograph of a set of 7,808 movie data collected. **b,** Workflow of Cryo-EM data processing of DDM1^HELLS^-nucleosome complex. Resolution estimation of final refinement is shown. **c,** Representation of selected 2D classes **d,** Angular distribution of particles used in final refinement. **e,** Fourier Shell Correlation curves for singly bound complex with a nominal resolution of 3.5 Å (FSC=0.143, broken lines)

**Extended Data Fig. 5.**
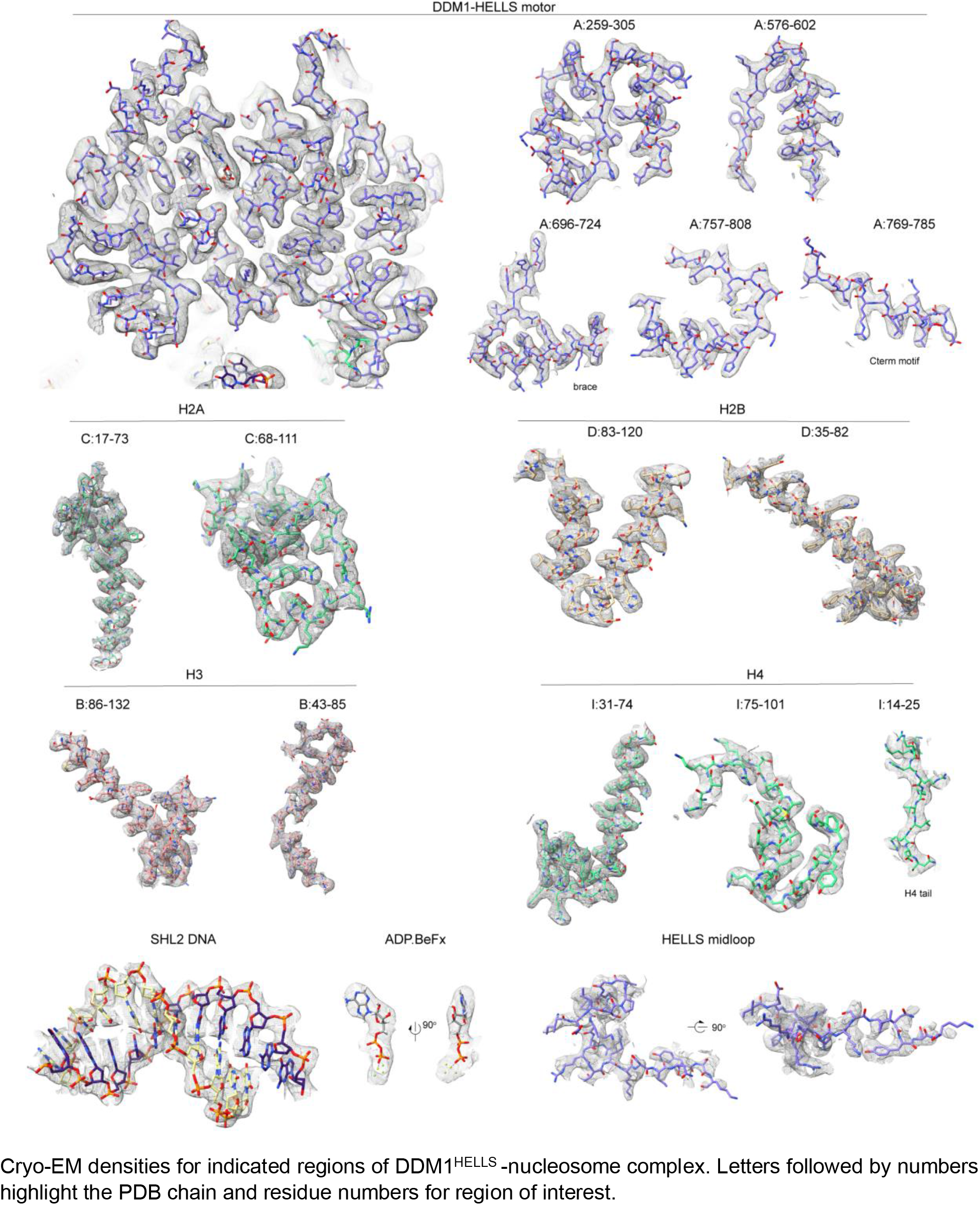
| Representative cryo-EM density maps of DDM1^HELLS^-nucleosome complex. Cryo-EM densities for indicated regions of DDM1^HELLS^-nucleosome complex. Letters followed by numbers highlight the PDB chain and residue numbers for region of interest.

**Extended Data Fig. 6.**
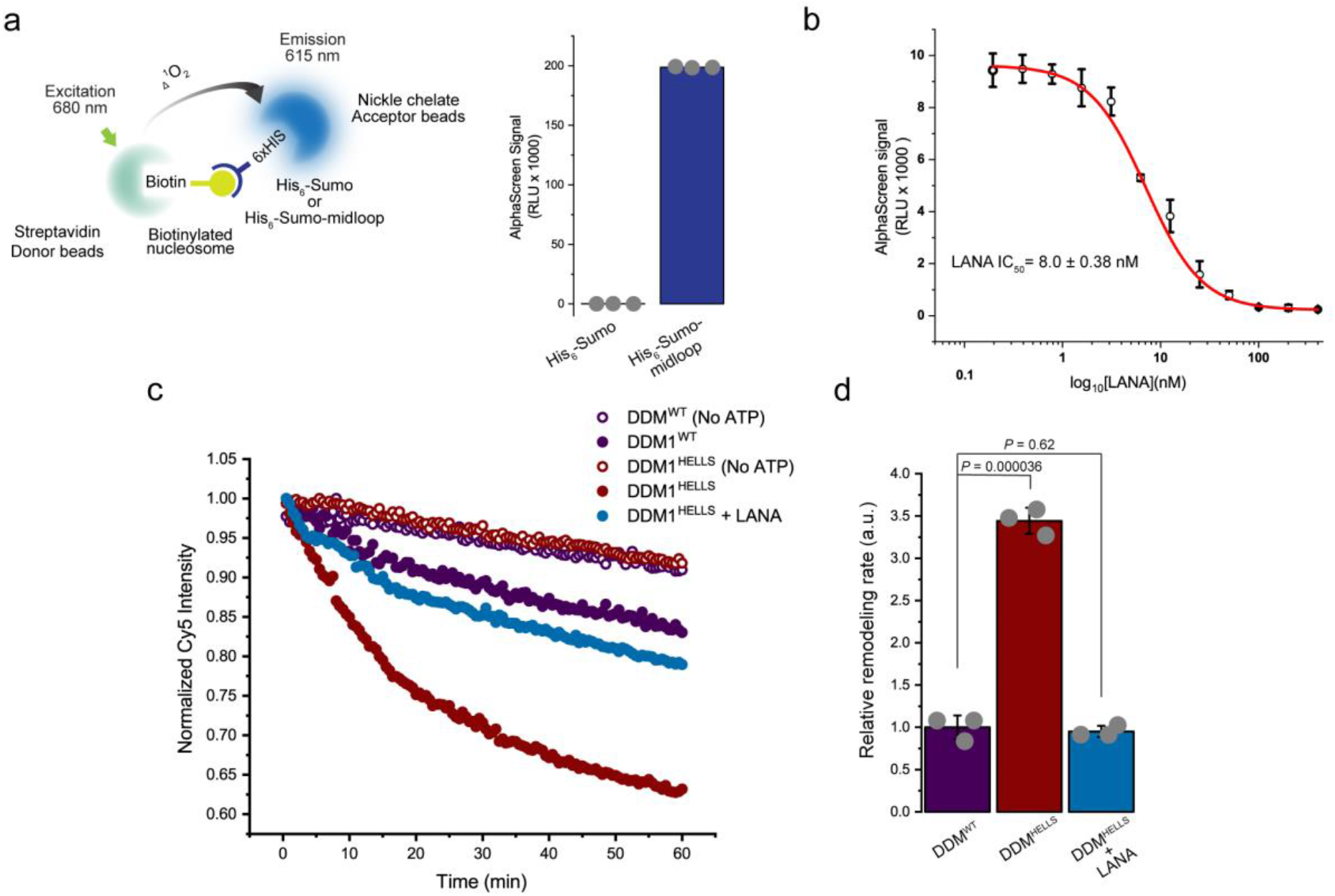
| HELLS midloop-nucleosome interactions. **a,** AlphaScreen experimental setup for midloop-nucleosome interaction. *(right)* Alphascreen signal acquired from SUMO-fused midloop-nucleosome interaction compared with SUMO only control. Data are mean ± s.d. (n=3 technical replicates). **b,** Competition assay of SUMO fused midloop-nucleosome interaction with LANA peptide. **c,** Representative FRET traces comparing DDM1 (1.0 µM) and DDM1^HELLS^ (1.0 µM) with and without addition of LANA peptide (100 µM) using 0.03 µM nucleosome substrate in the presence or absence of 2 mM ATP. **d**, Quantification of FRET traces in **c**. The error bars represent s.d. of three independent experiments. Statistical comparisons were performed with unpaired student’s two-tailed t-test.

**Extended Data Fig. 7.**
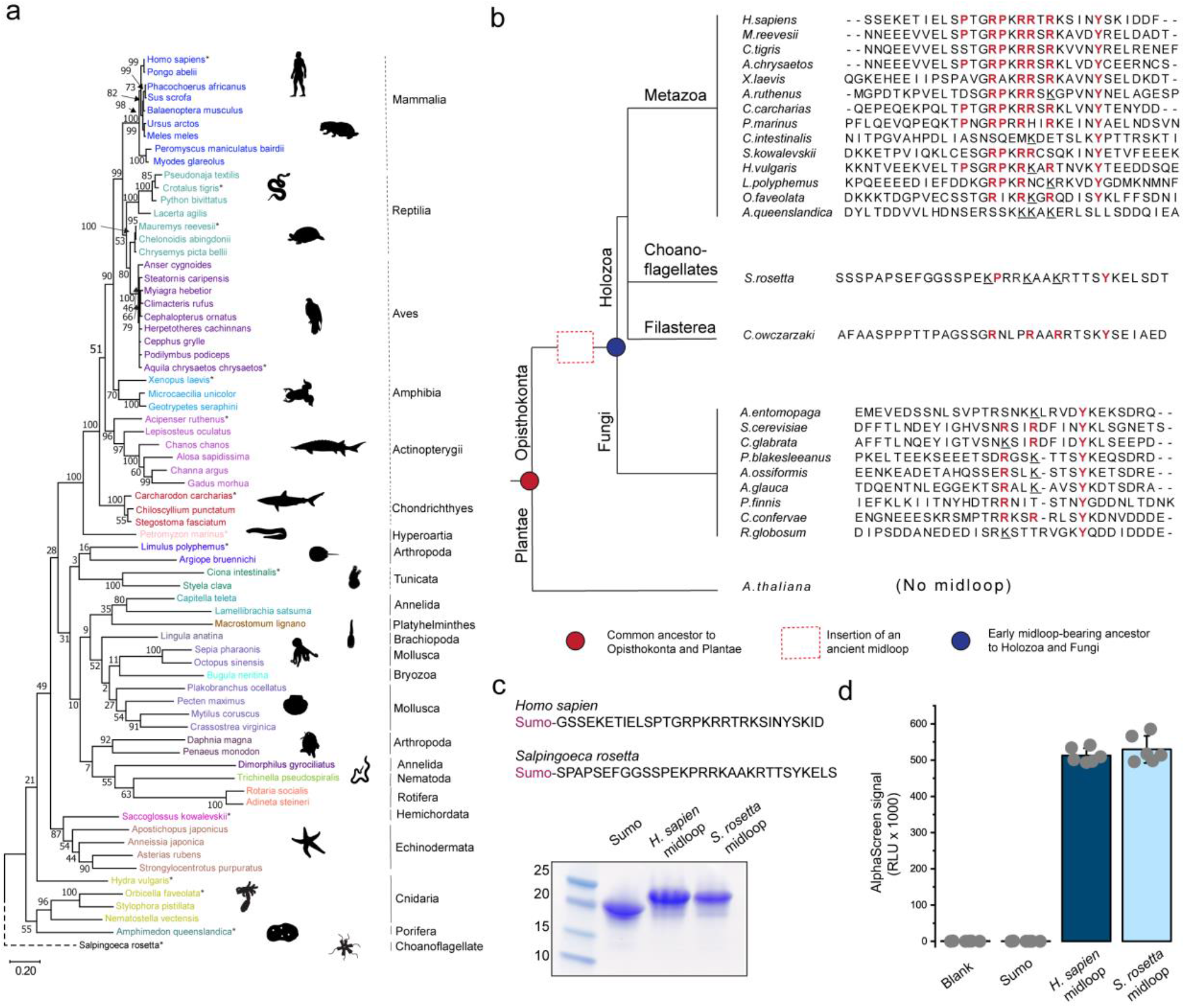
| Evolution of acidic-patch binding specificity of HELLS midloop. **a,** A phylogenetic tree of metazoan HELLS grouped and coloured by phylum (solid vertical line) or by class (broken vertical lines) for the phylum Chordata. Marked (asterisks) species were used to generate midloop alignment for **b**. Silhouettes of representative organisms were obtained from PhyloPic v2. The choanoflagellate, *Salpingoeca rosetta,* was used as the outgroup. Bootstrap statistics for 500 replicates are indicated on each node. All branch lengths are drawn to scale. **b,** Schematic model for the evolution of HELLS midloop (Not drawn to scale). Midloop sequence of the Holozoan supergroup was aligned with MUSCLE and positions of key residues are highlighted in bold red text based on alignment with *Homo sapiens* HELLS midloop. Lysine substitutions for arginine are underlined. Sequences from major fungi divisions are aligned and positions of potential interacting residues are marked based on our structure and *Homo sapiens* HELLS sequence. **c,** *Homo sapiens* and *Salpingoeca rosetta* HELLS midloop were tagged with SUMO and purified for Alphascreen assays. **d,** Alphascreen signals of His-tagged SUMO or SUMO- midloop fusion and biotin-tagged nucleosome. The error bars represent s.d. of six technical replicates.

**Extended Data Fig. 8.**
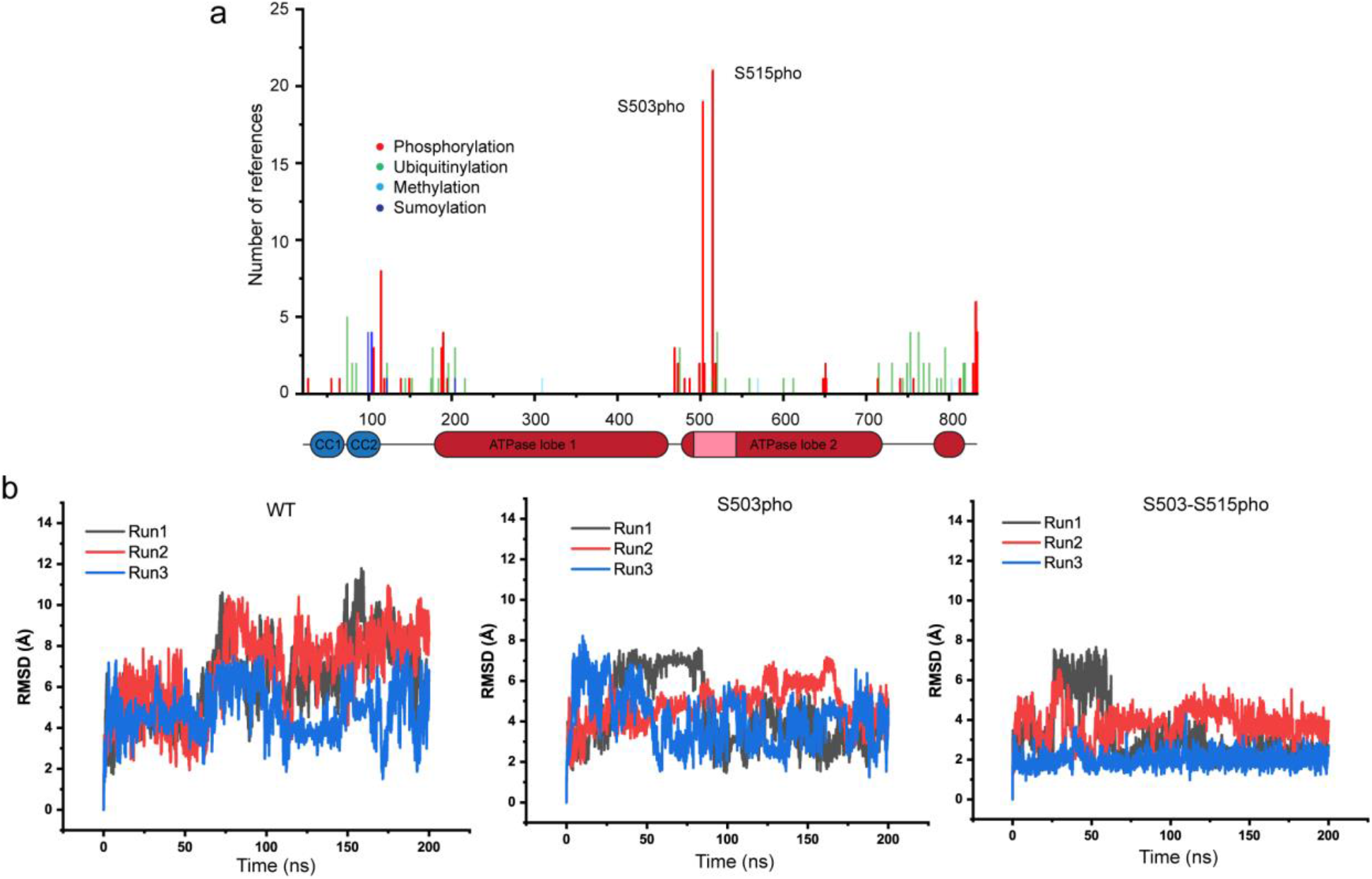
| Phosphorylation modulates conformational dynamics of HELLS midloop. **a,** A plot of the number of referenced post-translational modifications of HELLS curated from published literature (low and high-throughput experiments) in the PhosphoSitePlus database. **b**, Independent MD simulation for wild-type, S503p and S503p-S515p HELLS midloop.

**Extended Data Fig. 9.**
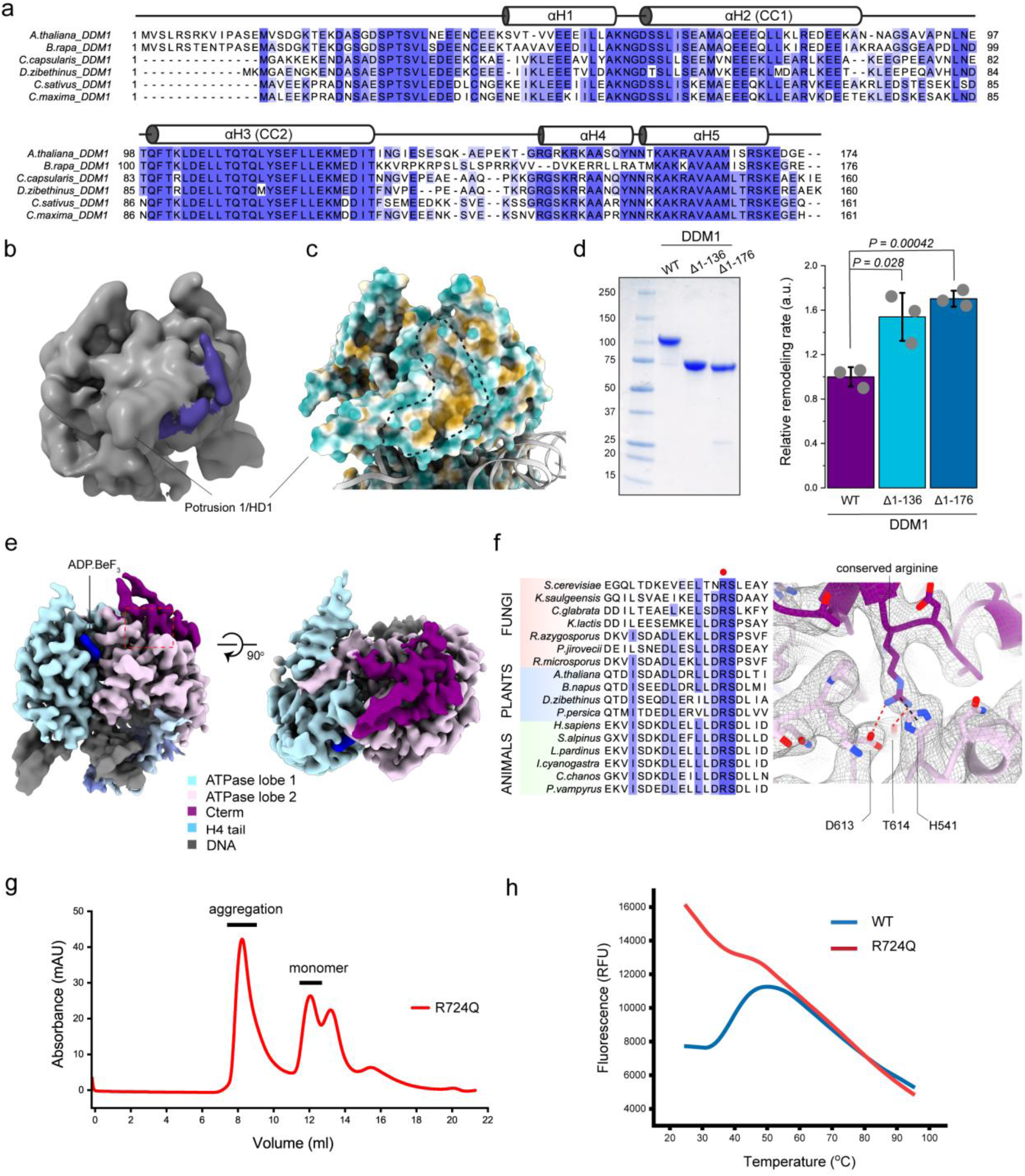
| Structural features of N- and C- terminus of DDM1. **a,** Sequence alignment of the N-terminal domain of DDM1 homologs. Predicted helices are shown above alignment. **b,** Low-pass filtered map revealing unmodelled N-terminus density (purple) packed against protrusion 1 of the DDM1 ATPase 1 motor. **c**, Surface representation of DDM1^motor^ coloured by hydrophobicity, with dotted lines marking the hydrophobic patch where the N-terminal helices pack. **d**, Coomassie-stained SDS-PAGE of DDM1 wild-type, DDM1 Δ1-136 and DDM1 Δ1-176. *Right,* Quantification of FRET traces comparing DDM1 wild-type and mutants. The error bars represent s.d. of three technical replicates. **e,** Two views of focused-refined DDM1^motor^ showing the packing of the C-terminus (magenta) on ATPase lobe 2 (pale magenta). ADP.BeF_x_ (blue) is shown bound to the active site. Red box highlights the Cterm motif. **f,** Sequence alignment of HELLS homologs from fungi (red), Plantae (blue) and metazoan (Green) revealing a highly conserved arginine residue. *Right,* the Cryo-EM density of the Cterm motif with atomic fitting reveals elaborate interaction of conserved arginine. **g**, Size-exclusion chromatography of R724Q mutant. **h**, Thermal denaturation profile of freeze-thawed monomeric DDM1^HELLS^ wild-type and R724Q mutant

**Extended Data Fig. 10.**
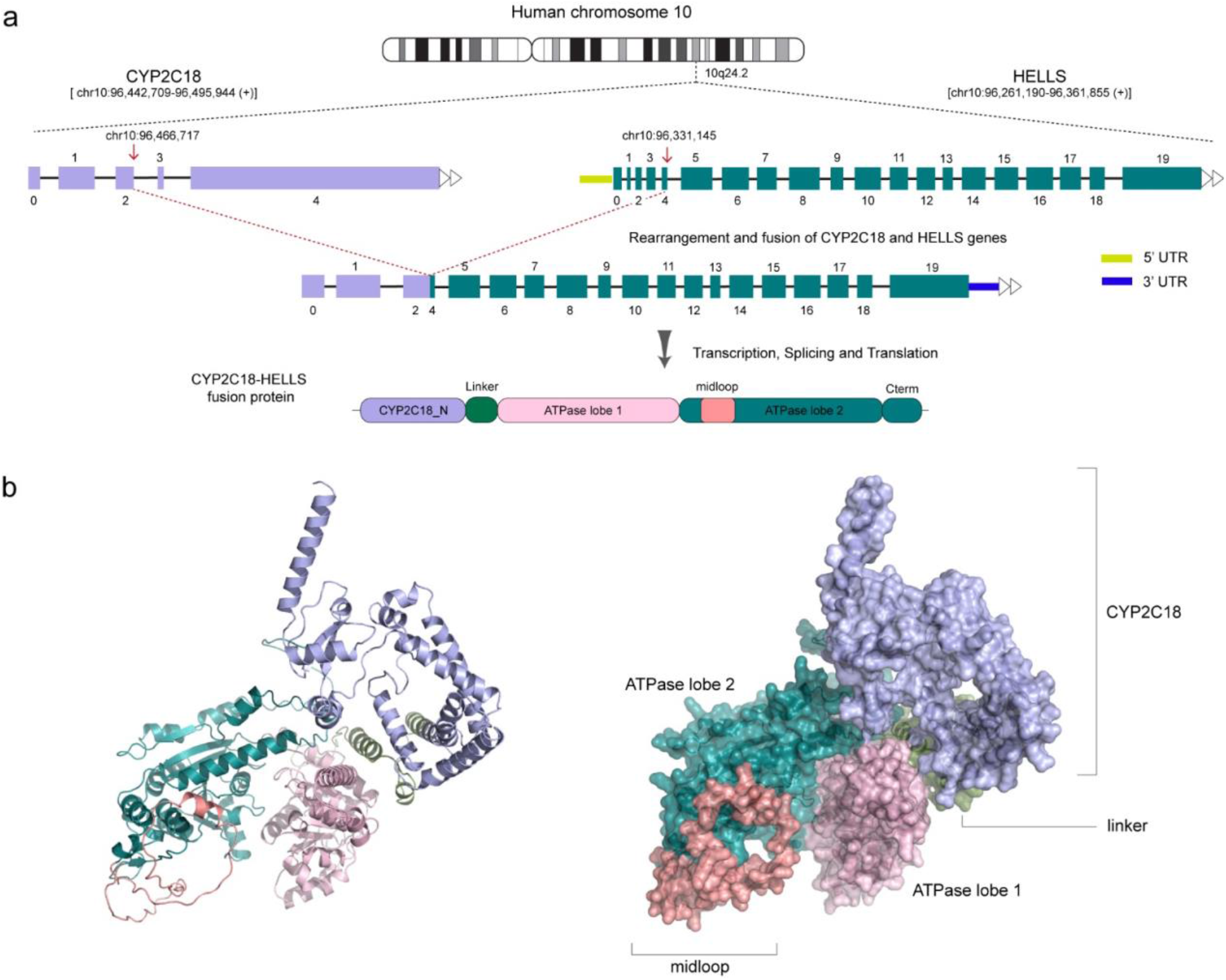
| Chromatin aberration creates a CYP2C18-HELLS fusion protein. **a,** Fusion of Exon 2 of CYP2C18 and Exon 4 of HELLS leads to formation of a chimera lacking the N- terminal regulatory domain of HELLS in a hepatocellular carcinoma tumour sample (TCGA-ZS-A9CF- 01A). Schematic representation is drawn based on annotations in the ChimerSeq database ^96^. **b**, Cartoon and surface representation of AlphaFold 2.0 prediction of the CYP2C18-HELLS fusion.

## ACKNOWLEDGEMENTS

We acknowledge the Arabidopsis Biological Resource Center at Ohio State University for providing cDNA for DDM1 (U24908). We would like to thank Dr. Claire Atkinson and the operating team of the High-Resolution Macromolecular cryo-Electron Microscopy facility (HRMEM) at UBC for their assistance in sample screening and data collection. HRMEM is funded by the Canadian Foundation of Innovation, BC Knowledge Development Fund, and the University of British Columbia.

## References

1. Clapier, C.R., Iwasa, J., Cairns, B.R. & Peterson, C.L. Mechanisms of action and regulation of ATP-dependent chromatin-remodelling complexes. Nature Reviews Molecular Cell Biology 18, 407–422 (2017).

2. Clapier, C.R. & Cairns, B.R. The Biology of Chromatin Remodeling Complexes. Annual Review of Biochemistry 78, 273–304 (2009).

3. Yan, L. & Chen, Z. A Unifying Mechanism of DNA Translocation Underlying Chromatin Remodeling. Trends in Biochemical Sciences 45, 217–227 (2020).

4. Nodelman, I.M. & Bowman, G.D. Biophysics of Chromatin Remodeling. Annual Review of Biophysics 50(2021).

5. Han, M. et al. A role for LSH in facilitating DNA methylation by DNMT1 through enhancing UHRF1 chromatin association. Nucleic Acids Research 48, 12116–12134 (2020).

6. Lyons, D.B. & Zilberman, D. DDM1 and Lsh remodelers allow methylation of DNA wrapped in nucleosomes. eLife 6(2017).

7. Termanis, A. et al. The SNF2 family ATPase LSH promotes cell-autonomousde novoDNA methylation in somatic cells. Nucleic Acids Research 44, 7592–7604 (2016).

8. Yan, Q., Cho, E., Lockett, S. & Muegge, K. Association of Lsh, a Regulator of DNA Methylation, with Pericentromeric Heterochromatin Is Dependent on Intact Heterochromatin. Molecular and Cellular Biology 23, 8416–8428 (2003).

9. Unoki, M., Funabiki, H., Velasco, G., Francastel, C. & Sasaki, H. CDCA7 and HELLS mutations undermine nonhomologous end joining in centromeric instability syndrome. Journal of Clinical Investigation 129, 78–92 (2018).

10. Jenness, C. et al. HELLS and CDCA7 comprise a bipartite nucleosome remodeling complex defective in ICF syndrome. Proceedings of the National Academy of Sciences 115, E876–E885 (2018).

11. Thijssen, P.E. et al. Mutations in CDCA7 and HELLS cause immunodeficiency–centromeric instability–facial anomalies syndrome. Nature Communications 6, 7870 (2015).

12. Shaked, H., Avivi-Ragolsky, N. & Levy, A.A. Involvement of the Arabidopsis SWI2/SNF2 Chromatin Remodeling Gene Family in DNA Damage Response and Recombination. Genetics 173, 985–994 (2006).

13. Qüesta, J.I., Fina, J.P. & Casati, P. DDM1 and ROS1 have a role in UV-B induced-and oxidative DNA damage in A. thaliana. Frontiers in Plant Science 4(2013).

14. Spruce, C. et al. HELLS and PRDM9 form a pioneer complex to open chromatin at meiotic recombination hot spots. Genes & Development 34, 398–412 (2020).

15. Basenko, E.Y., Kamei, M., Ji, L., Schmitz, R.J. & Lewis, Z.A. The LSH/DDM1 Homolog MUS-30 Is Required for Genome Stability, but Not for DNA Methylation in Neurospora crassa. PLOS Genetics 12, e1005790 (2016).

16. Burrage, J. et al. The SNF2 family ATPase LSH promotes phosphorylation of H2AX and efficient repair of DNA double-strand breaks in mammalian cells. Journal of Cell Science 125, 5524–5534 (2012).

17. Kollárovič, G., Topping, C.E., Shaw, E.P. & Chambers, A.L. The human HELLS chromatin remodelling protein promotes end resection to facilitate homologous recombination and contributes to DSB repair within heterochromatin. Nucleic Acids Research 48, 1872–1885 (2020).

18. Ni, K. & Muegge, K. LSH catalyzes ATP-driven exchange of histone variants macroH2A1 and macroH2A2. Nucleic Acids Research 49, 8024–8036 (2021).

19. Xu, X. et al. The epigenetic regulator LSH maintains fork protection and genomic stability via MacroH2A deposition and RAD51 filament formation. Nature Communications 12(2021).

20. Ni, K. et al. LSH mediates gene repression through macroH2A deposition. Nature Communications 11(2020).

21. Osakabe, A. et al. The chromatin remodeler DDM1 prevents transposon mobility through deposition of histone variant H2A.W. Nature Cell Biology 23, 391–400 (2021).

22. Imai, Y. et al. PRDM9 activity depends on HELLS and promotes local 5-hydroxymethylcytosine enrichment. eLife 9(2020).

23. Zhang, G. et al. Chromatin remodeler HELLS maintains glioma stem cells through E2F3 and MYC. JCI Insight 4(2019).

24. Jia, J. et al. Decrease in Lymphoid Specific Helicase and 5-hydroxymethylcytosine Is Associated with Metastasis and Genome Instability. Theranostics 7, 3920–3932 (2017).

25. De Dieuleveult, M. et al. The chromatin remodelling protein LSH/HELLS regulates the amount and distribution of DNA hydroxymethylation in the genome. Epigenetics 17, 422–443 (2022).

26. Chen, L. et al. DNA methylation modifier LSH inhibits p53 ubiquitination and transactivates p53 to promote lipid metabolism. Epigenetics & Chromatin 12(2019).

27. Myant, K. et al. LSH and G9a/GLP complex are required for developmentally programmed DNA methylation. Genome Research 21, 83–94 (2011).

28. Yang, R. et al. LSH interacts with and stabilizes GINS4 transcript that promotes tumourigenesis in non-small cell lung cancer. Journal of Experimental & Clinical Cancer Research 38(2019).

29. Wang, R. et al. The ratio of FoxA1 to FoxA2 in lung adenocarcinoma is regulated by LncRNA HOTAIR and chromatin remodeling factor LSH. Scientific Reports 5, 17826 (2016).

30. Benavente, C.A. et al. Chromatin remodelers HELLS and UHRF1 mediate the epigenetic deregulation of genes that drive retinoblastoma tumor progression. Oncotarget 5, 9594–9608 (2014).

31. Fragliasso, V. et al. The novel lncRNA BlackMamba controls the neoplastic phenotype of ALK− anaplastic large cell lymphoma by regulating the DNA helicase HELLS. Leukemia 34, 2964–2980 (2020).

32. Hou, X. et al. HELLS, a chromatin remodeler is highly expressed in pancreatic cancer and downregulation of it impairs tumor growth and sensitizes to cisplatin by reexpressing the tumor suppressor TGFBR3. Cancer Medicine (2020).

33. Waseem, A., Ali, M., Odell, E.W., Fortune, F. & Teh, M.-T. Downstream targets of FOXM1: CEP55 and HELLS are cancer progression markers of head and neck squamous cell carcinoma. Oral Oncology 46, 536–542 (2010).

34. Zocchi, L. et al. Chromatin remodeling protein HELLS is critical for retinoblastoma tumor initiation and progression. Oncogenesis 9(2020).

35. Robinson, M.H. et al. Upregulation of the chromatin remodeler HELLS is mediated by YAP1 in Sonic Hedgehog Medulloblastoma. Scientific Reports 9(2019).

36. Liu, X. et al. Downregulation of the Helicase Lymphoid-Specific (HELLS) Gene Impairs Cell Proliferation and Induces Cell Cycle Arrest in Colorectal Cancer Cells. OncoTargets and Therapy **Volume** 12, 10153–10163 (2019).

37. Von Eyss, B. et al. The SNF2-like helicase HELLS mediates E2F3-dependent transcription and cellular transformation. The EMBO Journal 31, 972–985 (2012).

38. Han, Y. et al. Lsh/HELLS regulates self-renewal/proliferation of neural stem/progenitor cells. Scientific Reports 7 (2017).

39. He, X. et al. Chromatin Remodeling Factor LSH Drives Cancer Progression by Suppressing the Activity of Fumarate Hydratase. Cancer Research 76, 5743–5755 (2016).

40. Liu, S. & Tao, Y.-G. Chromatin remodeling factor LSH affects fumarate hydratase as a cancer driver. Chinese Journal of Cancer 35(2016).

41. Liu, N. et al. The cross-talk between methylation and phosphorylation in lymphoid-specific helicase drives cancer stem-like properties. Signal Transduction and Targeted Therapy 5(2020).

42. Law, C.T. et al. HELLS Regulates Chromatin Remodeling and Epigenetic Silencing of Multiple Tumor Suppressor Genes in Human Hepatocellular Carcinoma. Hepatology 69, 2013–2030 (2019).

43. Brzeski, J. & Jerzmanowski, A. Deficient in DNA Methylation 1 (DDM1) Defines a Novel Family of Chromatin-remodeling Factors. Journal of Biological Chemistry 278, 823–828 (2003).

44. Hauk, G., Mcknight, J.N., Nodelman, I.M. & Bowman, G.D. The Chromodomains of the Chd1 Chromatin Remodeler Regulate DNA Access to the ATPase Motor. Molecular Cell 39, 711–723 (2010).

45. Clapier, C.R. & Cairns, B.R. Regulation of ISWI involves inhibitory modules antagonized by nucleosomal epitopes. Nature 492, 280–284 (2012).

46. Paul, S. & Bartholomew, B. Regulation of ATP-dependent chromatin remodelers: accelerators/brakes, anchors and sensors. Biochemical Society Transactions 46, 1423–1430 (2018).

47. Shahian, T. & Narlikar, G.J. Analysis of Changes in Nucleosome Conformation Using Fluorescence Resonance Energy Transfer. in Methods in Molecular Biology 337–349 (Humana Press, 2012).

48. Li, M. et al. Mechanism of DNA translocation underlying chromatin remodelling by Snf2. Nature 567, 409–413 (2019).

49. Thomä, N.H. et al. Structure of the SWI2/SNF2 chromatin-remodeling domain of eukaryotic Rad54. Nature Structural & Molecular Biology 12, 350–356 (2005).

50. Flaus, A. Identification of multiple distinct Snf2 subfamilies with conserved structural motifs. Nucleic Acids Research 34, 2887–2905 (2006).

51. Wang, L., Chen, K. & Chen, Z. Structural basis of ALC1/CHD1L autoinhibition and the mechanism of activation by the nucleosome. Nature Communications 12(2021).

52. Armache, J.P. et al. Cryo-EM structures of remodeler-nucleosome intermediates suggest allosteric control through the nucleosome. eLife 8(2019).

53. Farnung, L., Ochmann, M. & Cramer, P. Nucleosome-CHD4 chromatin remodeler structure maps human disease mutations. eLife 9(2020).

54. Nodelman, I.M. et al. Nucleosome recognition and DNA distortion by the Chd1 remodeler in a nucleotide-free state. Nature Structural & Molecular Biology 29, 121–129 (2022).

55. Clapier, C.R. A critical epitope for substrate recognition by the nucleosome remodeling ATPase ISWI. Nucleic Acids Research 30, 649–655 (2002).

56. Hwang, W.L., Deindl, S., Harada, B.T. & Zhuang, X. Histone H4 tail mediates allosteric regulation of nucleosome remodelling by linker DNA. Nature 512, 213–217 (2014).

57. Yan, L., Wang, L., Tian, Y., Xia, X. & Chen, Z. Structure and regulation of the chromatin remodeller ISWI. Nature 540, 466–469 (2016).

58. Clapier, C.R., LäNgst, G., Corona, D.F.V., Becker, P.B. & Nightingale, K.P. Critical Role for the Histone H4 N Terminus in Nucleosome Remodeling by ISWI. Molecular and Cellular Biology 21, 875–883 (2001).

59. Barbera, A.J. et al. The nucleosomal surface as a docking station for Kaposi’s sarcoma herpesvirus LANA. Science 311, 856–61 (2006).

60. McGinty, R.K., Tan, Song. Principles of nucleosome recognition by chromatin factors and enzymes. CURRENT OPINION IN STRUCTURAL BIOLOGY 71, 16–26 (2021).

61. Yuan, J., Chen, K., Zhang, W. & Chen, Z. Structure of human chromatin-remodelling PBAF complex bound to a nucleosome. Nature 605, 166–171 (2022).

62. Sen, P., Ghosh, S., Pugh, B.F. & Bartholomew, B. A new, highly conserved domain in Swi2/Snf2 is required for SWI/SNF remodeling. Nucleic Acids Research 39, 9155–9166 (2011).

63. Fairclough, S.R. et al. Premetazoan genome evolution and the regulation of cell differentiation in the choanoflagellate Salpingoeca rosetta. Genome Biol 14, R15 (2013).

64. Yan, L., Wu, H., Li, X., Gao, N. & Chen, Z. Structures of the ISWI–nucleosome complex reveal a conserved mechanism of chromatin remodeling. Nature Structural & Molecular Biology 26, 258–266 (2019).

65. Lehmann, L.C. et al. Mechanistic Insights into Autoinhibition of the Oncogenic Chromatin Remodeler ALC1. Molecular Cell 68, 847–859.e7 (2017).

66. Wang, L. et al. Regulation of the Rhp26 ERCC6/CSB chromatin remodeler by a novel conserved leucine latch motif. Proceedings of the National Academy of Sciences 111, 18566–18571 (2014).

67. Wang, W. et al. Molecular basis of chromatin remodeling by Rhp26, a yeast CSB ortholog. Proceedings of the National Academy of Sciences 116, 6120–6129 (2019).

68. Ferreira, H., Flaus, A. & Owen-Hughes, T. Histone Modifications Influence the Action of Snf2 Family Remodelling Enzymes by Different Mechanisms. Journal of Molecular Biology 374, 563–579 (2007).

69. Shogren-Knaak, M. et al. Histone H4-K16 acetylation controls chromatin structure and protein interactions. Science 311, 844–7 (2006).

70. Corcoran, E.T. et al. Systematic histone H4 replacement in Arabidopsis thaliana reveals a role for H4R17 in regulating flowering time. The Plant Cell (2022).

71. Zhang, X. et al. Cis-and trans-regulation by histone H4 basic patch R17/R19 in metazoan development. Open Biology 12(2022).

72. Lehmann, L.C. et al. Mechanistic Insights into Regulation of the ALC1 Remodeler by the Nucleosome Acidic Patch. Cell Reports 33, 108529 (2020).

73. Gamarra, N., Johnson, S.L., Trnka, M.J., Burlingame, A.L. & Narlikar, G.J. The nucleosomal acidic patch relieves auto-inhibition by the ISWI remodeler SNF2h. eLife 7(2018).

74. Gioacchini, N. & Peterson, C.L. SWR1C catalyzes H2A.Z deposition by coupling ATPase activity to the nucleosome acidic patch. (Cold Spring Harbor Laboratory, 2021).

75. Xia, X., Liu, X., Li, T., Fang, X. & Chen, Z. Structure of chromatin remodeler Swi2/Snf2 in the resting state. Nature Structural & Molecular Biology 23, 722–729 (2016).

76. Zegzouti, H., Zdanovskaia, M., Hsiao, K. & Goueli, S.A. ADP-Glo: A Bioluminescent and homogeneous ADP monitoring assay for kinases. Assay Drug Dev Technol 7, 560–72 (2009).

77. Kiianitsa, K., Solinger, J.A. & Heyer, W.-D. NADH-coupled microplate photometric assay for kinetic studies of ATP-hydrolyzing enzymes with low and high specific activities. Analytical Biochemistry 321, 266–271 (2003).

78. Luger, K., Rechsteiner, T.J. & Richmond, T.J. Expression and Purification of Recombinant Histones and Nucleosome Reconstitution. in Chromatin Protocols 1–16 (Humana Press).

79. Dao, H.T., Dul, B.E., Dann, G.P., Liszczak, G.P. & Muir, T.W. A basic motif anchoring ISWI to nucleosome acidic patch regulates nucleosome spacing. Nature Chemical Biology 16, 134–142 (2020).

80. Waterhouse, A.M., Procter, J.B., Martin, D.M.A., Clamp, M. & Barton, G.J. Jalview Version 2--a multiple sequence alignment editor and analysis workbench. Bioinformatics 25, 1189–1191 (2009).

81. Tamura, K., Stecher, G. & Kumar, S. MEGA11: Molecular Evolutionary Genetics Analysis Version 11. Molecular Biology and Evolution 38, 3022–3027 (2021).

82. Jones, D.T., Taylor, W.R. & Thornton, J.M. The rapid generation of mutation data matrices from protein sequences. Bioinformatics 8, 275–282 (1992).

83. Felsenstein, J. Confidence Limits on Phylogenies: An Approach Using the Bootstrap. Evolution 39, 783 (1985).

84. Jo, S., Kim, T., Iyer, V.G. & Im, W. CHARMM-GUI: A web-based graphical user interface for CHARMM. Journal of Computational Chemistry 29, 1859–1865 (2008).

85. Huang, J. et al. CHARMM36m: an improved force field for folded and intrinsically disordered proteins. Nature Methods 14, 71–73 (2017).

86. Hess, B., Bekker, H., Berendsen, H.J.C. & Fraaije, J.G.E.M. LINCS: A linear constraint solver for molecular simulations. Journal of Computational Chemistry 18, 1463–1472 (1997).

87. Van Der Spoel, D. et al. GROMACS: fast, flexible, and free. J Comput Chem 26, 1701–18 (2005).

88. Huynh, K. & Partch, C.L. Analysis of Protein Stability and Ligand Interactions by Thermal Shift Assay. Current Protocols in Protein Science 79(2015).

89. Punjani, A., Rubinstein, J.L., Fleet, D.J. & Brubaker, M.A. cryoSPARC: algorithms for rapid unsupervised cryo-EM structure determination. Nature Methods 14, 290–296 (2017).

90. Jumper, J. et al. Highly accurate protein structure prediction with AlphaFold. Nature 596, 583–589 (2021).

91. Pettersen, E.F. et al. UCSF ChimeraX: Structure visualization for researchers, educators, and developers. Protein Science 30, 70–82 (2021).

92. Croll, T.I. ISOLDE: a physically realistic environment for model building into low-resolution electron-density maps. Acta Crystallographica Section D Structural Biology 74, 519–530 (2018).

93. Emsley, P., Lohkamp, B., Scott, W.G. & Cowtan, K. Features and development of Coot. Acta Crystallographica Section D Biological Crystallography 66, 486–501 (2010).

94. Liebschner, D. et al. Macromolecular structure determination using X-rays, neutrons and electrons: recent developments in Phenix. Acta Crystallographica Section D Structural Biology 75, 861–877 (2019).

95. Chen, V.B., et al. MolProbity: all-atom structure validation for macromolecular crystallography. Acta Crystallographica Section D Biological Crystallography 66, 12–21 (2010).

96. Jang, Y.E. et al. ChimerDB 4.0: an updated and expanded database of fusion genes. Nucleic Acids Research (2019).

