## Supplementary figures for "Cryo-EM structure of DDM1-HELLS chimera bound to nucleosome reveals a mechanism of chromatin remodeling and disease regulation"

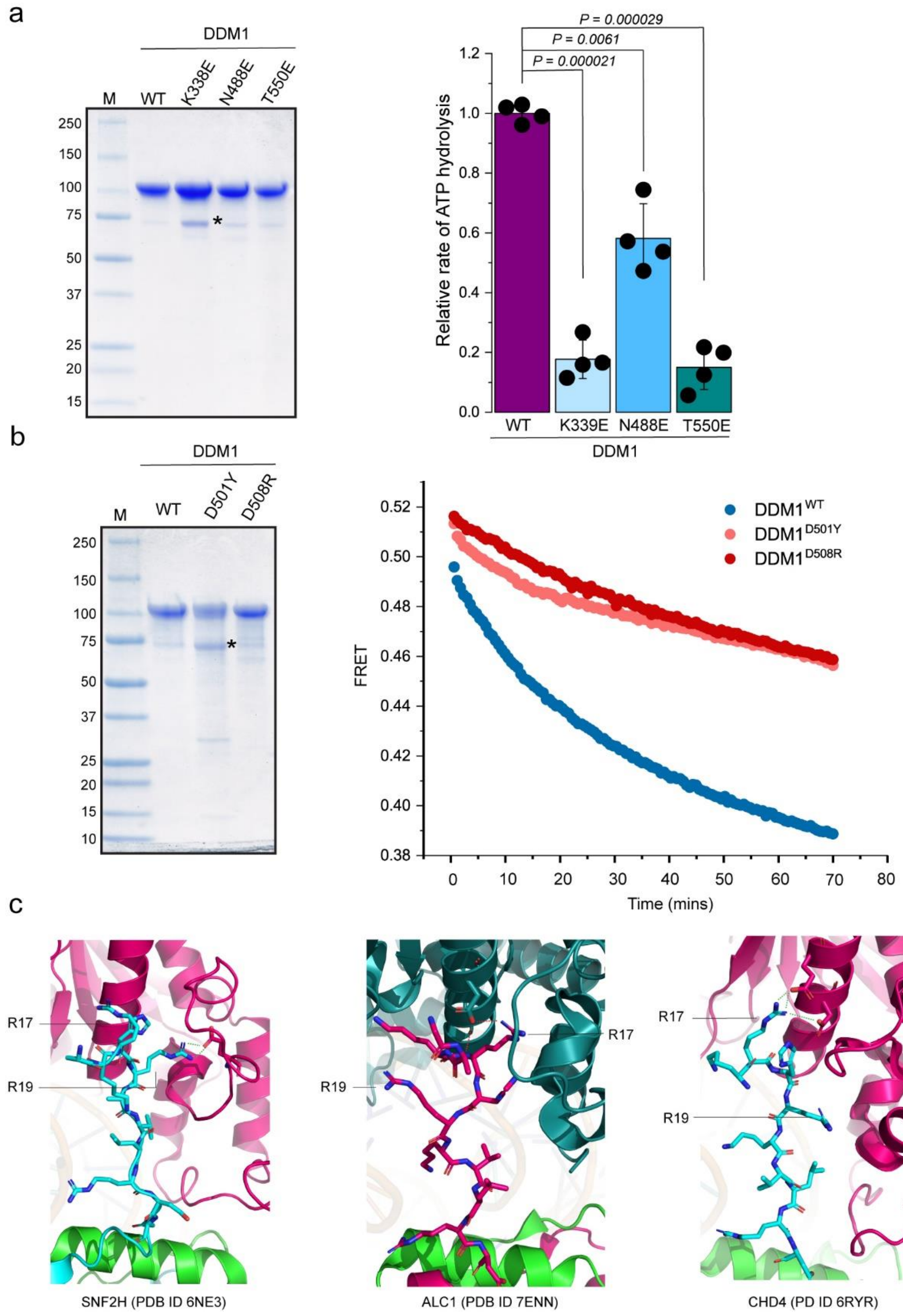


**Supplementary Fig. 1 | Comparison of DDM1 wild-type and mutant. a,** *(left)* Coomassie-stained SDS-PAGE of purified DDM1 and DNA binding mutants. M is protein standards *(right)* NADH-coupled DNA (99-mer random sequence) stimulated ATPase activity of DDM1 and mutant proteins. **b**, *(left)* Coomassie-stained SDS-PAGE of purified DDM1 and histone binding mutants. *(right)* Representative FRET traces used to generate **Fig. 4c**. **c,** Comparism of different mode of H4 N-terminal tail-motor interactions of snf2 remodellors

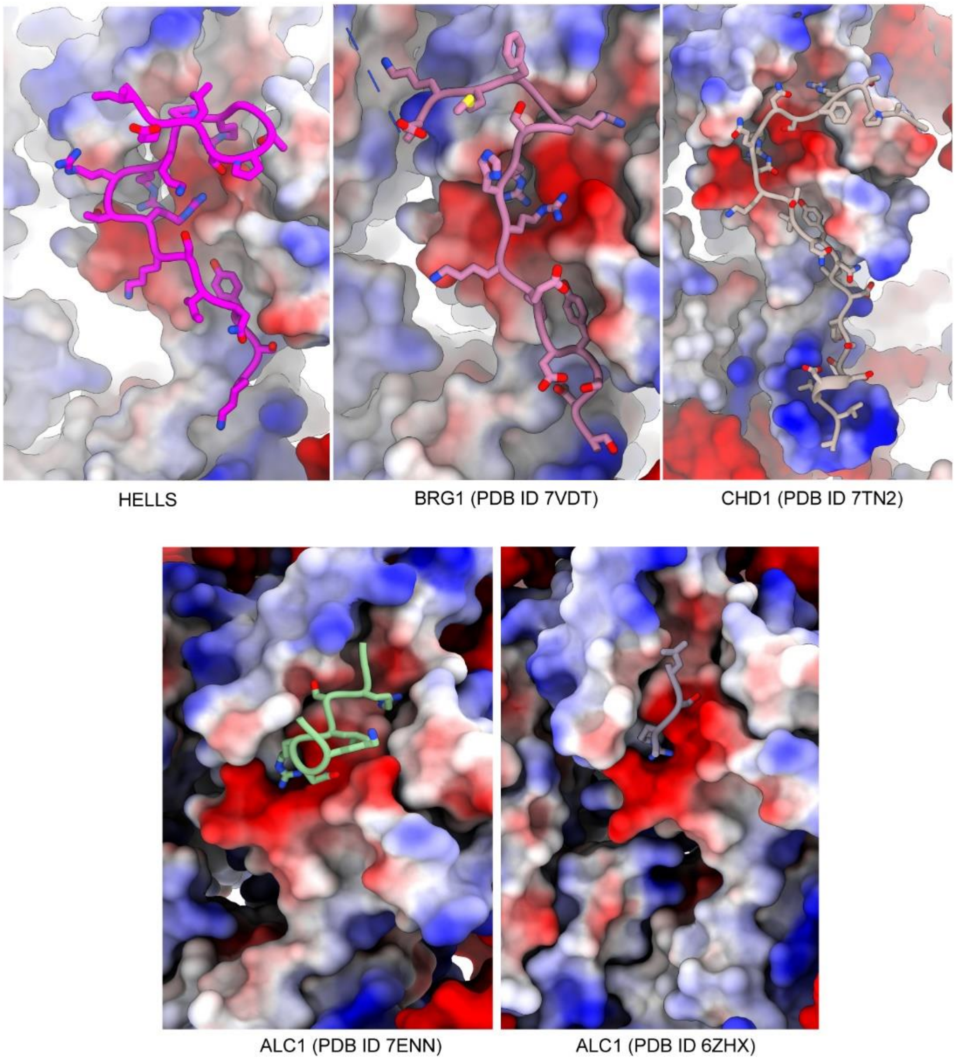


**Supplementary Fig. 2 | Comparison of interactions between Snf2 remodellers and nucleosome acidic patch**

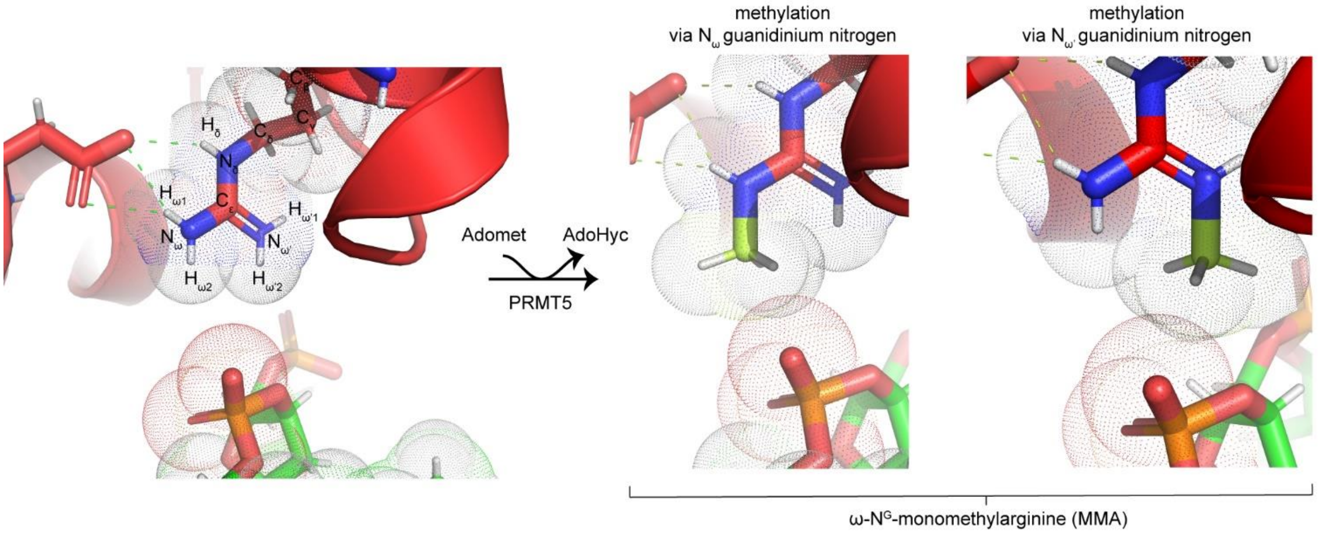


**Supplementary Fig. 3 | *In silico* methylation of R288 (HELLS R309).** Methylation was performed within the Maestro software suit (Schrödinger) and van der Waals volume visualized in pymol (Schrödinger).

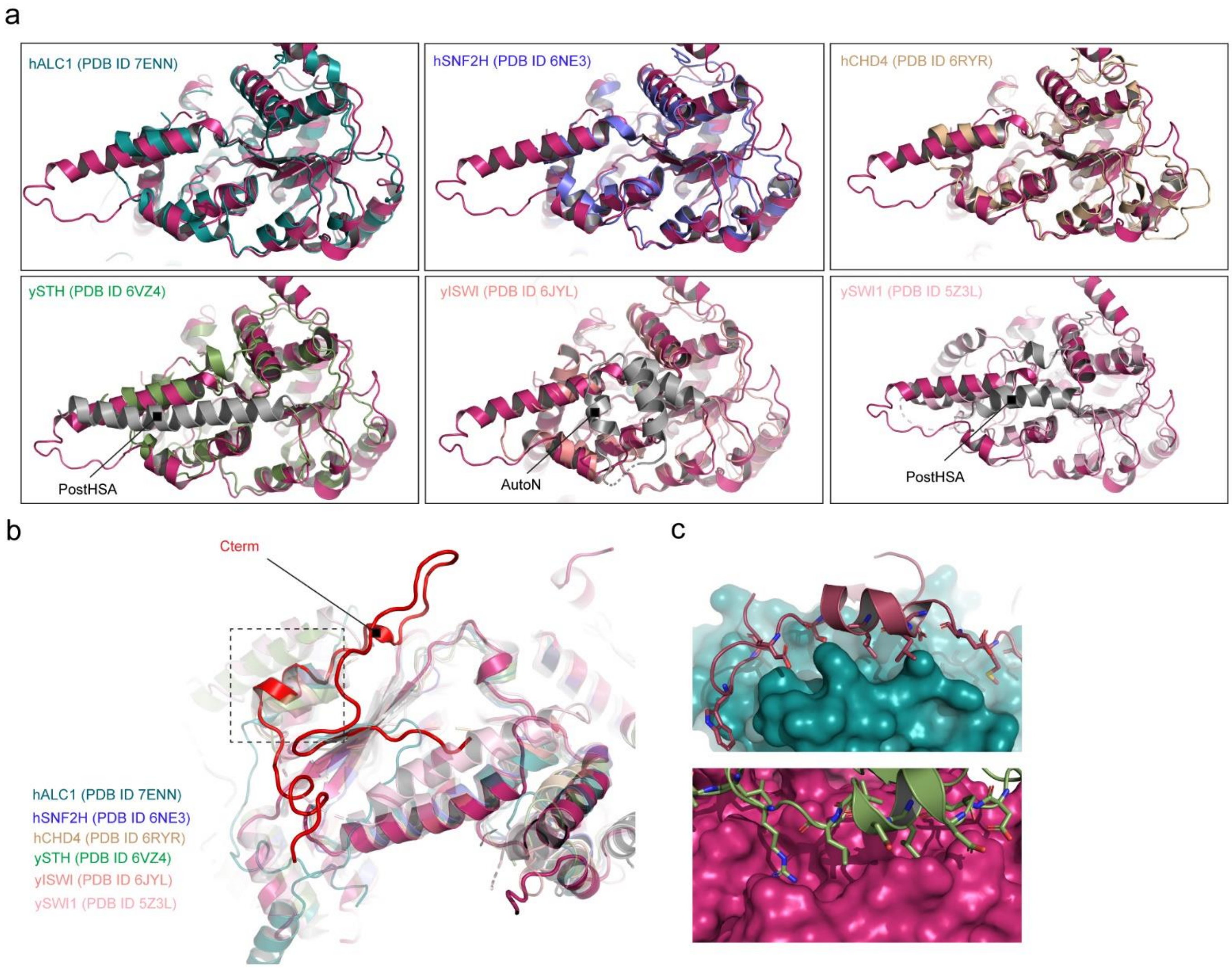


**Supplementary Fig. 4. Comparism of motor domain of DDM1 and other snf2 motor. a,** Comparism of the potrusion I region of DDM1 (*deep pink*) and other remodellors. **b**, C-terminus comparism of DDM1 (*red*) and othe snf2 motors. **c**, Comparism of the mode of interaction of the Cterm motif of DDM1 (*below*) and a corresponding region of hALC1 (*above*)

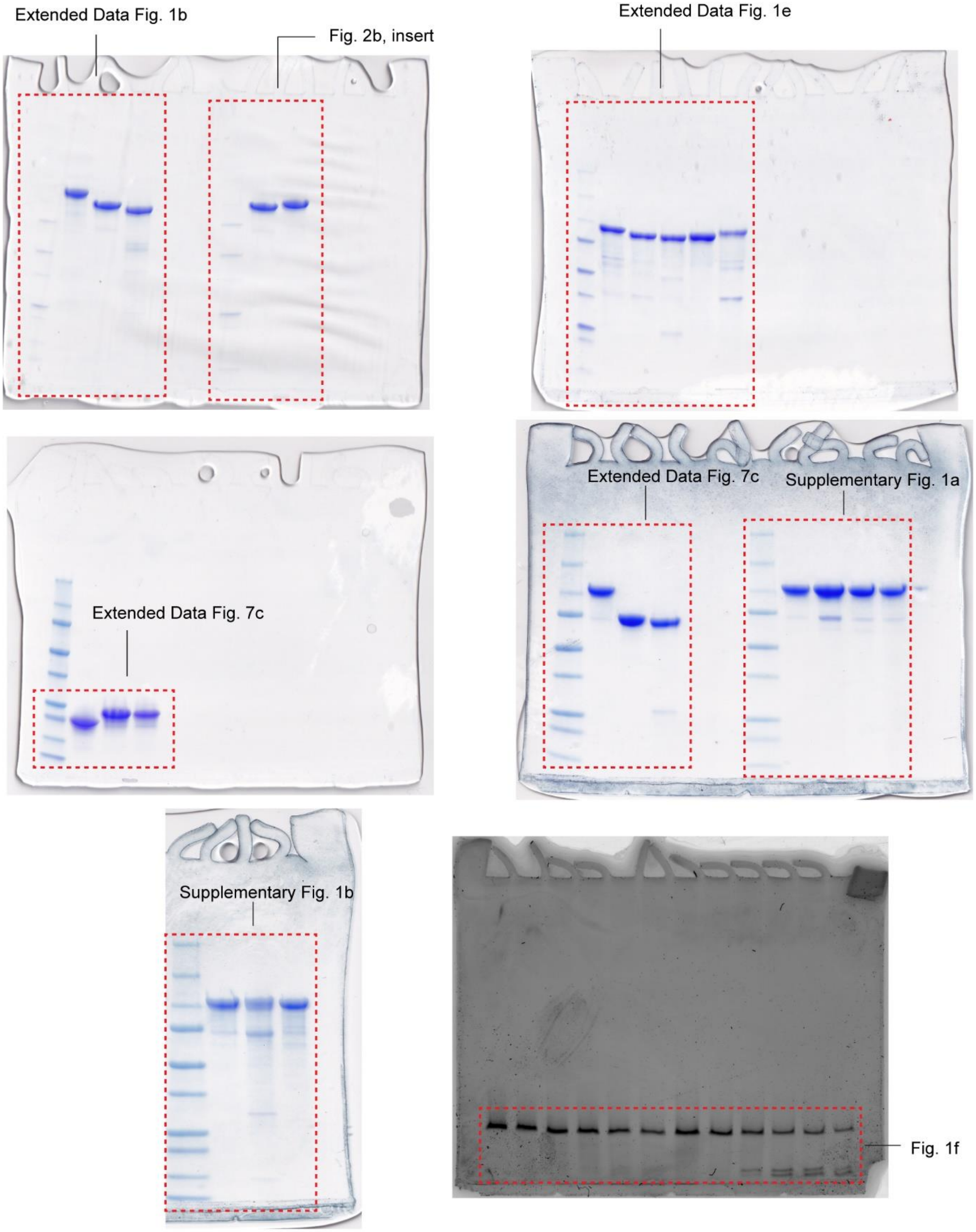


**Supplementary Fig. 5. Uncropped gels used for the preparation of corresponding figures**
